# Integrative genomics approach identifies conserved transcriptomic networks in Alzheimer’s disease

**DOI:** 10.1101/695221

**Authors:** Samuel Morabito, Emily Miyoshi, Neethu Michael, Vivek Swarup

**Affiliations:** Mathematical, Computational and Systems Biology (MCSB) Program, University of California, Irvine, CA 92697, USA; Department of Neurobiology and Behavior, University of California, Irvine, CA 92697, USA; Institute for Memory Impairments and Neurological Disorders (MIND), University of California, Irvine, CA 92697, USA

## Abstract

Alzheimer’s disease (AD) is a devastating neurological disorder characterized by changes in cell-type proportions and consequently marked alterations of the transcriptome. Here we use a data-driven systems biology approach across multiple cohorts of human AD, encompassing different brain regions, and integrate with multi-scale datasets comprising of DNA methylation, histone acetylation, transcriptome- and genome-wide association studies as well as quantitative trait loci to define the genetic architecture of AD. We perform co-expression network analysis across more than twelve hundred human brain samples, identifying robust AD-associated dysregulation of the transcriptome, unaltered in normal human aging. We further integrate co-expression modules with single-cell transcriptome generated from 27,321 nuclei from postmortem human brain to identify AD-specific transcriptional changes and assess cell-type proportion changes in the human AD brain. We also show that genetic variants of AD are enriched in a glial AD-associated module and identify key transcription factors regulating co-expressed modules. Additionally, we validate our results in multiple published human AD datasets which are easily accessible using our online resource (https://swaruplab.bio.uci.edu/consensusAD).

## Introduction

Alzheimer’s disease (AD) is a progressive neurodegenerative disease, which is characterized by impairment in cognitive and executive functions, including memory loss.^1, 2^ AD is pathologically characterized by the presence of neurofibrillary tangles and senile plaques, which are comprised of hyperphosphorylated microtubule associated protein tau (*MAPT*; tau) inclusions and amyloid beta (Aβ) deposits, respectively.^2, 3^ While aging is the major non-genetic risk factor for AD, several genetic risk factors, including mutations in MAPT, have been found and associated with the disease.^4–7^ However despite the identification of causal genetic mutations associated with AD and genes implicated in genome-wide association studies (GWAS) of the disease,^6, 7^ the mechanisms of neurodegeneration are still poorly understood and are complex, impeding the design of therapeutic intervention.

Transcriptomic approaches coupled with gene network analysis are powerful in investigating quantitative molecular phenotypes and pathways underlying disease progression in a genome-wide manner. These approaches have the potential to not only unravel new pathways underlying disease mechanisms, but also to identify disease regulators, which themselves can be likely targets for drug development. Within this context, the National Institute of Aging (NIA) has developed a Target Identification and Preclinical Validation Project of the Accelerating Medicines Project – Alzheimer’s Disease (AMP-AD) consortium,^8^ whose goal is to integrate high-throughput genomic and molecular data from the diseased brain within a network driven structure.^8, 9^ Under this program, several AD-specific large-scale RNA sequencing (RNA-seq) projects have been conducted. These projects identified transcriptomic networks and splicing events that are altered in the cortex of the AD brain and evaluated the potential role of a viral transcriptome in modifying disease pathology.^9–16^ However, these studies were limited to specific brain banks, and large meta-analysis across hundreds of samples with integrative genomic approaches is pertinent to robustly identify disease-specific signatures in an unbiased manner.

Here we have taken a functional genomics and integrative systems biology approach to identify human AD-specific transcriptomic alterations conserved across multiple cohorts and multiple cortical regions. We applied a consensus network analysis approach and identified robust disease relevant co-expression modules, thus simultaneously analyzing more than 1200 postmortem brain samples across brain banks, making this a unique study. We found several neuronal and glial modules that are altered in AD but not in normal human aging. While the neuronal processes were related to synaptic transmission, proteasomal degradation and mitochondrial dysfunction, the glial processes were enriched in inflammation, immune pathways and metabolism. We also found that one of the glial modules is enriched in AD candidate risk genes from AD GWAS datasets. We used generic and brain-specific ATAC-seq and ChIP-seq datasets to define potential transcription factors regulating the neuronal and glial modules. In addition, we have combined various orthogonal datasets including DNA methylation, histone acetylation (H3K9ac), transcriptome and genome-wide association studies (TWAS and GWAS) as well as expression and methylation quantitative loci (eQTLs and mQTLs) and integrated with co-expression network analysis to provide a comprehensive picture of genetic alterations associated with AD.

Massive changes in cell composition are concurrent with the progression of the disease, impairing accurate estimation of transcriptomic changes from bulk tissue data. To circumvent this problem, we generated single-nuclei transcriptomics data from normal post-mortem human brain and used the obtained transcriptomic signatures to accurately estimate cell-type proportion changes from bulk RNA-seq samples. Finally, we validated the findings in multiple orthogonal published datasets of AD encompassing different brain regions, highlighting the robustness of our findings. Together, our findings define a core set of transcriptomic changes altered during the progression of the disease and highlight several regulators of these changes, which may be used as therapeutic targets.

## Results

In order to identify key transcriptional changes in AD, we analyzed RNA-seq data from three different studies—Mayo Clinic Brain Bank (Mayo),^10^ the Religious Orders Study and Memory and Aging Project (ROSMAP),^14^ and Mount Sinai School of Medicine (MSSM).^17^ The Mayo dataset included temporal cortex samples (n = 160),^10^ while the ROSMAP dataset was composed of prefrontal cortex samples (n = 632).^14^ The MSSM cohort included samples from four different regions—parahippocampal gyrus, inferior frontal gyrus, superior temporal gyrus, and the frontal pole (n = 476, Figure 1).^17^ Altogether, we used these datasets to examine disease-related changes that were conserved across more than a thousand individuals from different cortical regions. All datasets were uniformly processed and used for differential gene expression analysis and consensus weighted gene co-expression network analysis (see Supplemental Table 1 for all case information).

**Figure 1.**
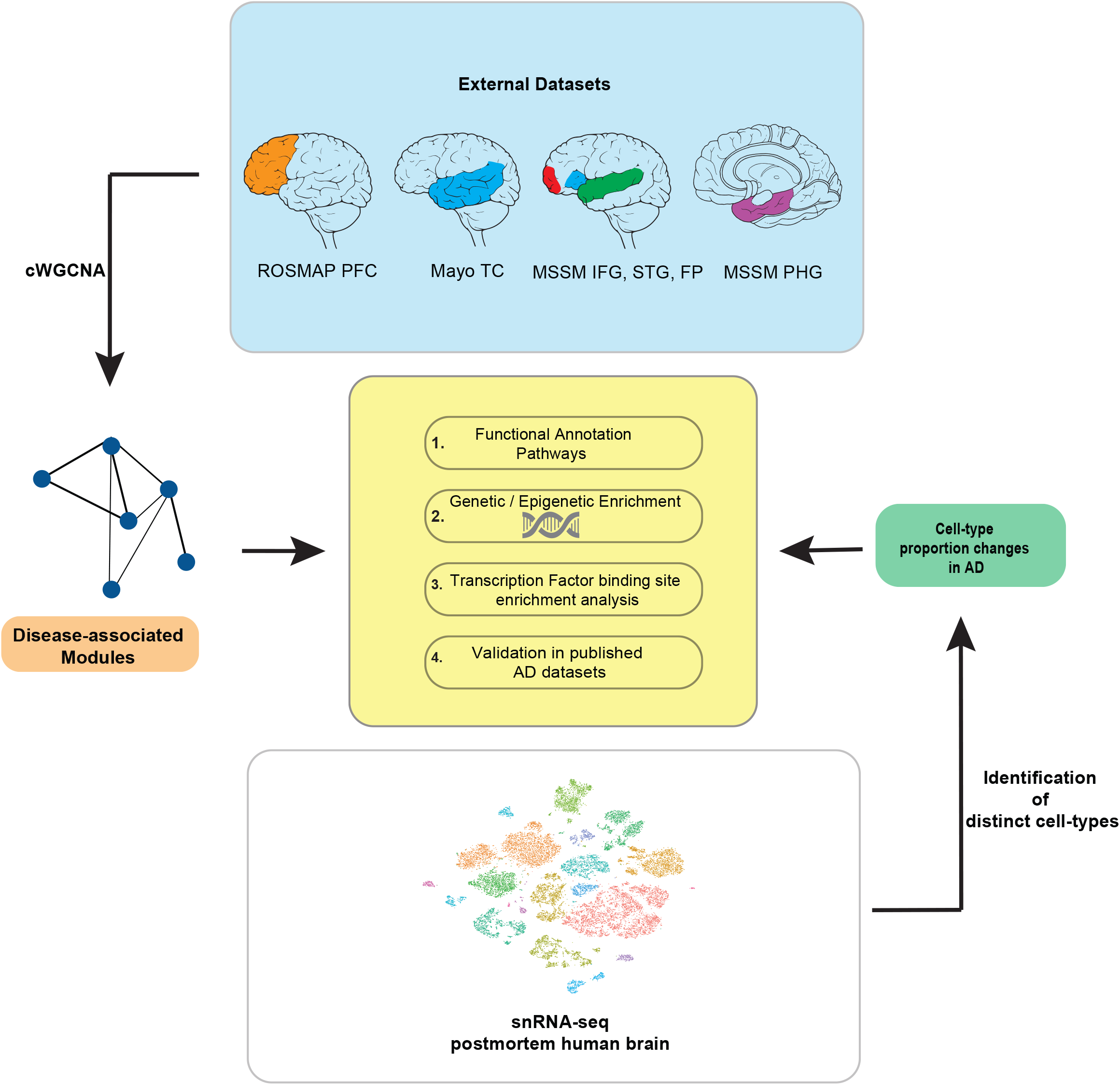
Schematic representation of the various analyses performed in the study Consensus weighted gene co-expression network analysis (cWGCNA) with datasets from three different studies—Mayo Clinic Brain Bank (Mayo) temporal cortex (TC); Religious Orders Study and Memory and Aging Project (ROSMAP) prefrontal cortex (PFC); Mount Sinai School of Medicine (MSSM) para-hippocampal gyrus (PHG), inferior frontal gyrus (IFG), superior temporal gyrus (STG), and frontal pole (FP)—identified Alzheimer’s disease (AD)-associated modules. Distinct cell-types were detected using single-nuclei RNA-sequencing (snRNA-seq) from postmortem human brain tissue revealing cell-type proportion changes in AD. Disease-associated modules and cell-type clusters were used for functional pathway analysis in addition to GWAS enrichment, transcription factor binding site enrichment and validation in published datasets.

### Transcriptional alterations of protein-coding and non-coding genes in AD

We focused our analyses on a subset of the Mayo temporal cortex dataset in which AD and control individuals were age-matched (Discovery Set, n = 104, Supplemental Table 1) and found protein-coding genes differentially expressed in AD (Figure 2a, Supplemental Table 2). Gene ontology (GO) analysis revealed enrichment of terms related to immune and defense response in upregulated protein-coding genes and terms related to synaptic transmission in downregulated genes (Figure 2b). In addition, the upregulated genes were enriched in endothelial, astrocytic, and microglial marker genes, while the downregulated were enriched in neuronal genes (Fig 2c). We then confirmed that our differential expression analysis of protein-coding genes was preserved in the remainder of the Mayo cohort (Replication Set, Figure 2d) and other studies (Figure 2e-g, Supplemental Figure 1a-g).

**Figure 2.**
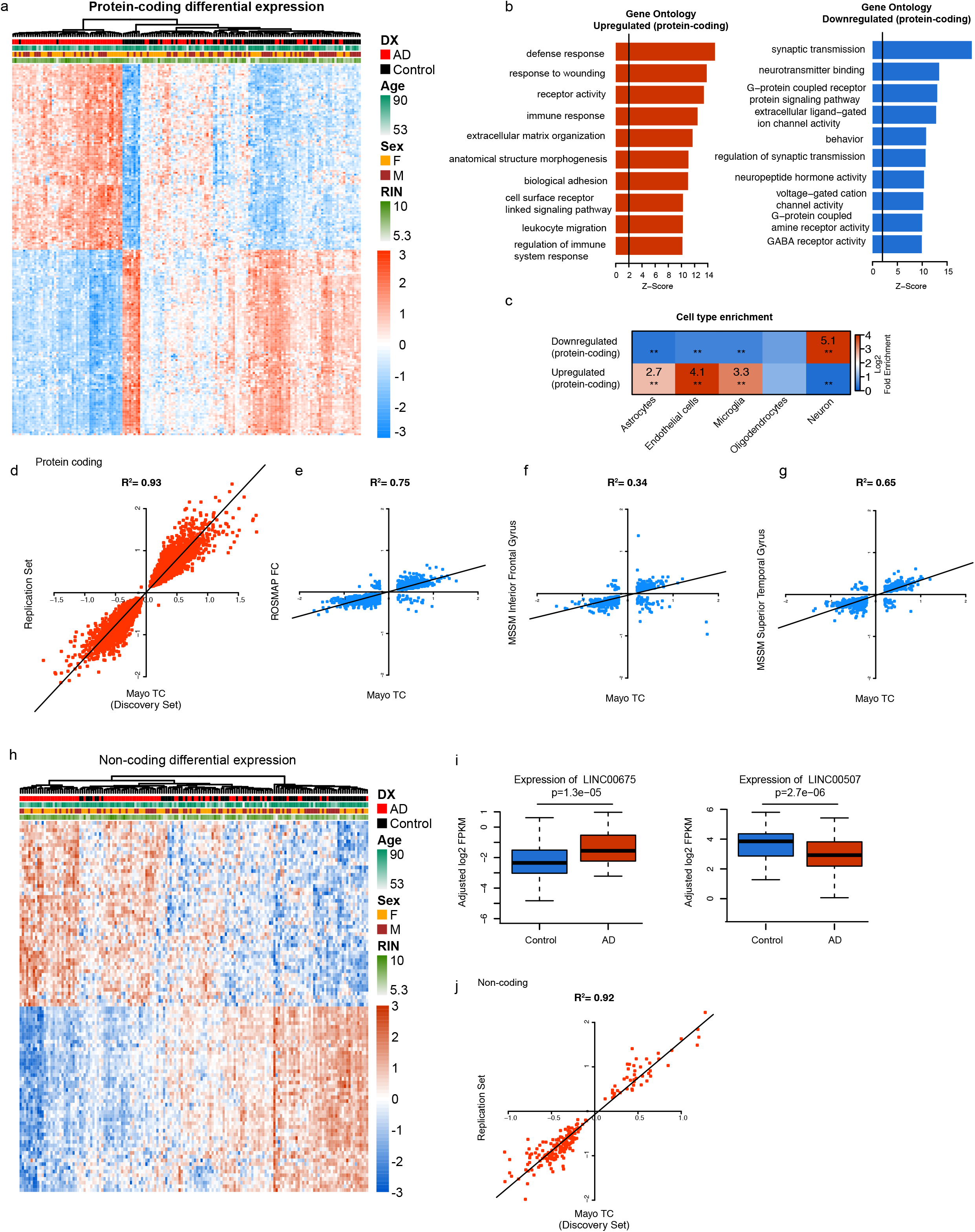
Differential expression analysis in protein-coding and non-coding RNA (**a**) Heat map of differentially expressed protein-coding genes identified in Mayo TC dataset with covariates diagnosis (DX), age, sex, and RNA integrity number (RIN). (**b**) Gene ontology term enrichment of the upregulated (left) and downregulated (right) protein-coding genes. (**c**) Enrichment of upregulated and downregulated protein-coding genes in different cell-types. (**d**) Correlation of differentially expressed protein-coding genes from Mayo TC (Discovery Set) with Mayo Replication Set. (**e-g**) Correlation of Mayo TC with ROSMAP FC (**e**), MSSM inferior frontal gyrus (**f**), and MSSM superior temporal gyrus (**g**). (**h**) Heat map of differentially expressed protein-coding genes identified in Mayo TC with covariates diagnosis (DX), age, sex, and RIN. (**i**) Boxplots showing change in expression of LINC00675 (left) and LINC00507 (right) with diagnosis. (**j**) Correlation of differentially expressed non-coding genes from Mayo Discovery Set with Mayo Replication Set. Sample sizes are mentioned in Supplemental Table 1.

Further, we discovered differential expression of non-coding genes with disease (Figure 2h) and identified long non-coding RNAs (lncRNAs) significantly upregulated or downregulated in AD (Figure 2i, Supplemental Table 2). lncRNAs play gene regulatory roles, and dysregulation of lncRNAs has been implicated in various diseases including AD.^18, 19^ LINC00675, which we found upregulated in AD, previously has been associated with cervical and colorectal cancer, impacting cell proliferation and migration.^20, 21^ On the other hand, we found LINC00507 expression decreased with disease state, and LINC00507 has been shown to be expressed specifically in the cortex and in an age-dependent manner.^22^ However, it is still unclear what is the functional significance of LINC00507, which may be important for understanding its role in disease. Since the other cohorts sequenced only mRNA, they could not be used for comparison, but we again confirmed our results in the Replication Set (Figure 2j).

### Identification of cell-type clusters in the adult human brain

Because the brain’s cell-type composition changes over the course of AD, gene expression results from RNA-seq performed on bulk tissue tend to reflect these major cell-type changes, likely causing us to overlook biologically relevant changes because they appear insignificant. Single-cell RNA-seq allows us to resolve transcriptional changes by individual cell-types, but it is currently not practical for large cohorts. However, we can use single-cell RNA-seq in order to better inform our analyses from bulk tissue RNA-seq. Using postmortem human brain tissue from neurologically healthy controls (n = 5, Supplemental Table 1), we isolated nuclei and performed single-nuclei RNA-seq (snRNA-seq, n = 27,321 nuclei) to clarify cell-type proportions and the corresponding transcriptional profiles of the adult human cortex. Through unsupervised clustering, we identified 18 transcriptionally distinct populations, visualized by t-SNE plot (Figure 3a, Supplemental Table 3). We found both neuronal and non-neuronal subpopulations: 7 excitatory neuron, 4 inhibitory neuron, 2 astrocyte, and 2 oligodendrocyte subpopulations. We additionally found clusters consistent with endothelial cells, microglia, and oligodendrocyte precursor cells (OPCs), and there was one cluster we were unable to confirm its cell-type origin. The majority of the isolated nuclei was classified as excitatory neurons (Figure 3b).

**Figure 3.**
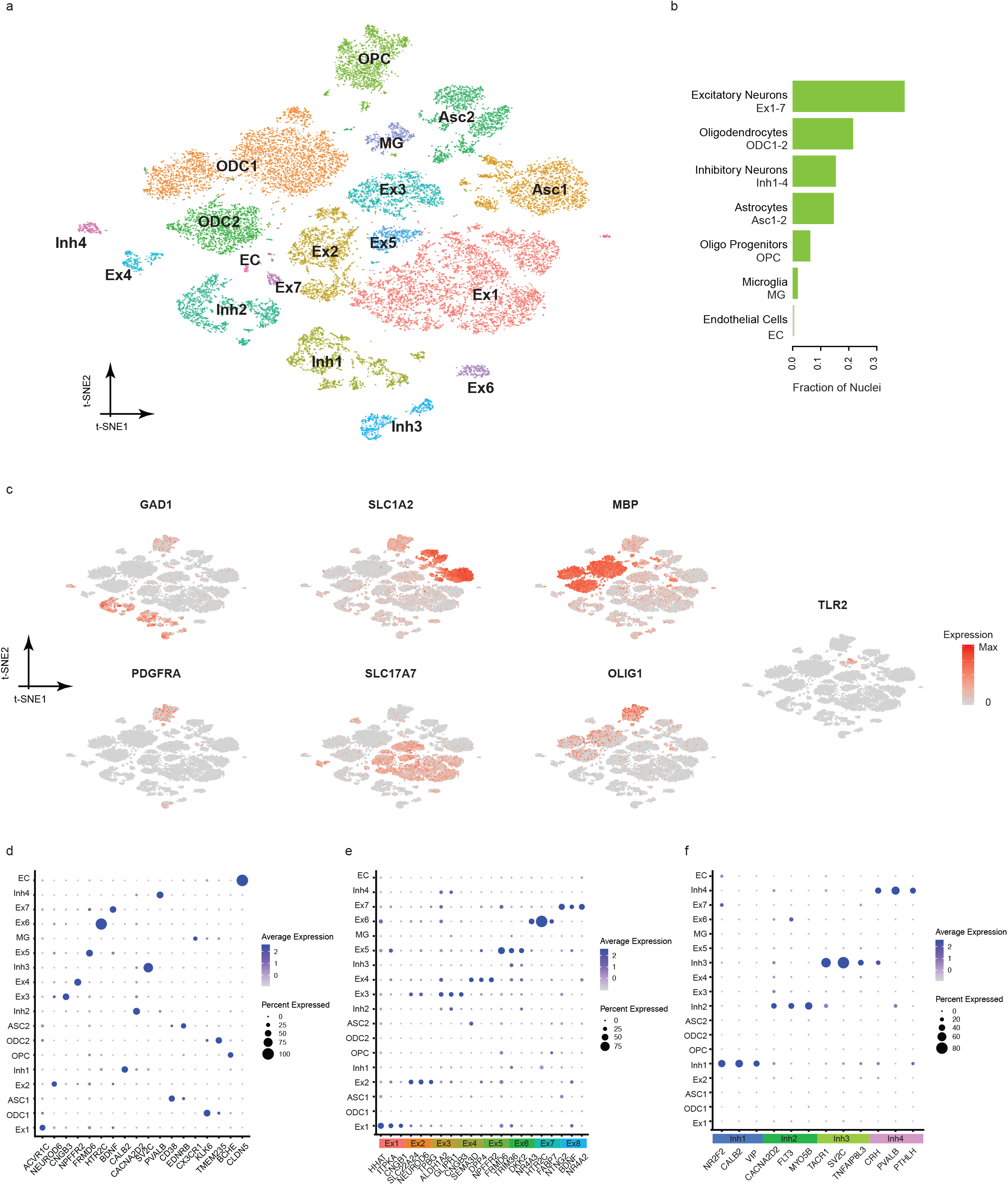
Single nuclei RNA-seq identifies robust cell-clusters in adult human brain (**a**) Cell-type clusters visualized with t-SNE plot. Cell-types were labeled *post hoc* based on marker gene enrichment. (**b**) Bar graph indicating fraction of nuclei identified as each cell-type. (**c**) Marker gene enrichment in cell-type clusters visualized with same t-SNE plot as in (**a**). (**d-f**) Dot plot showing differentially expressed cell-type markers (**d**), excitatory neuronal markers (**e**), and inhibitory neuronal markers (**f**) in cell-type clusters. Color denotes average gene expression, while size denotes percent of nuclei expressing the gene.

We classified our cell-type clusters based on enrichment of known cell-type markers (Figure 3c) and published single-cell clusters.^23, 24^ We identified gene markers specific to each cell-type cluster (Figure 3d, Supplemental Figure 2a). These included previously established cell-type markers, such as PVALB, NEUROD6, and CX3CR1, and layer-specific markers like SV2C and HTR2C.^24, 25^ Additionally, we utilized the Allen Human Brain Atlas to verify the expression of these marker genes in the human cortex (Supplemental Figure 2b). Because we found multiple neuronal subpopulations, we wanted to characterize them further and identified three selectively and highly co-expressed genes in each neuronal cluster (Figure 3e-f). We were able to identify our excitatory neuron clusters 7 and 8 as deep layer neurons, while our inhibitory neuron cluster 3 likely originates from layer 3 neurons. In addition, we found inhibitory neuron subpopulations similar to Darmanis et al.^26^—our inhibitory neuron clusters 1 and 4, CALB2-VIP and CRH-PVALB neurons, respectively.

### Cell-type proportion changes in AD

We then reasoned that our cell-type clusters from snRNA-seq could be leveraged to examine changes in cell-type proportions in AD across different brain regions. AD progresses in an anatomical manner,^1, 27^ which has suggested that pathology spread relies on synapses;^28^ however, in the context of transcriptomic studies, gene expression changes may be dependent on these region-specific cellular changes. Further, large bulk RNA-seq datasets may be affected due to differences in tissue sampling. We devised an algorithm in order to estimate the cell-type composition in bulk tissue RNA-seq data using our cell-type clusters (see Methods). Using the specificity score obtained from single-cell data, we independently estimated cell-type composition in each of the bulk RNA-seq datasets used. We found, in the Mayo Discovery Set, all excitatory and inhibitory neuronal subpopulations decreased with AD, as expected (Figure 4a-b), and non-neuronal cell-types increased, with the exception of OPCs (Figure 4c-e). We found similar patterns in the other datasets, but we noticed RNA-seq datasets from temporal cortical regions demonstrated more pronounced changes in cell-type proportions than frontal cortical regions (Supplemental Figure 3-4). This regional difference resembles the progression of pathology in which the temporal cortex is affected before the frontal cortex in AD.^1, 27^

**Figure 4.**
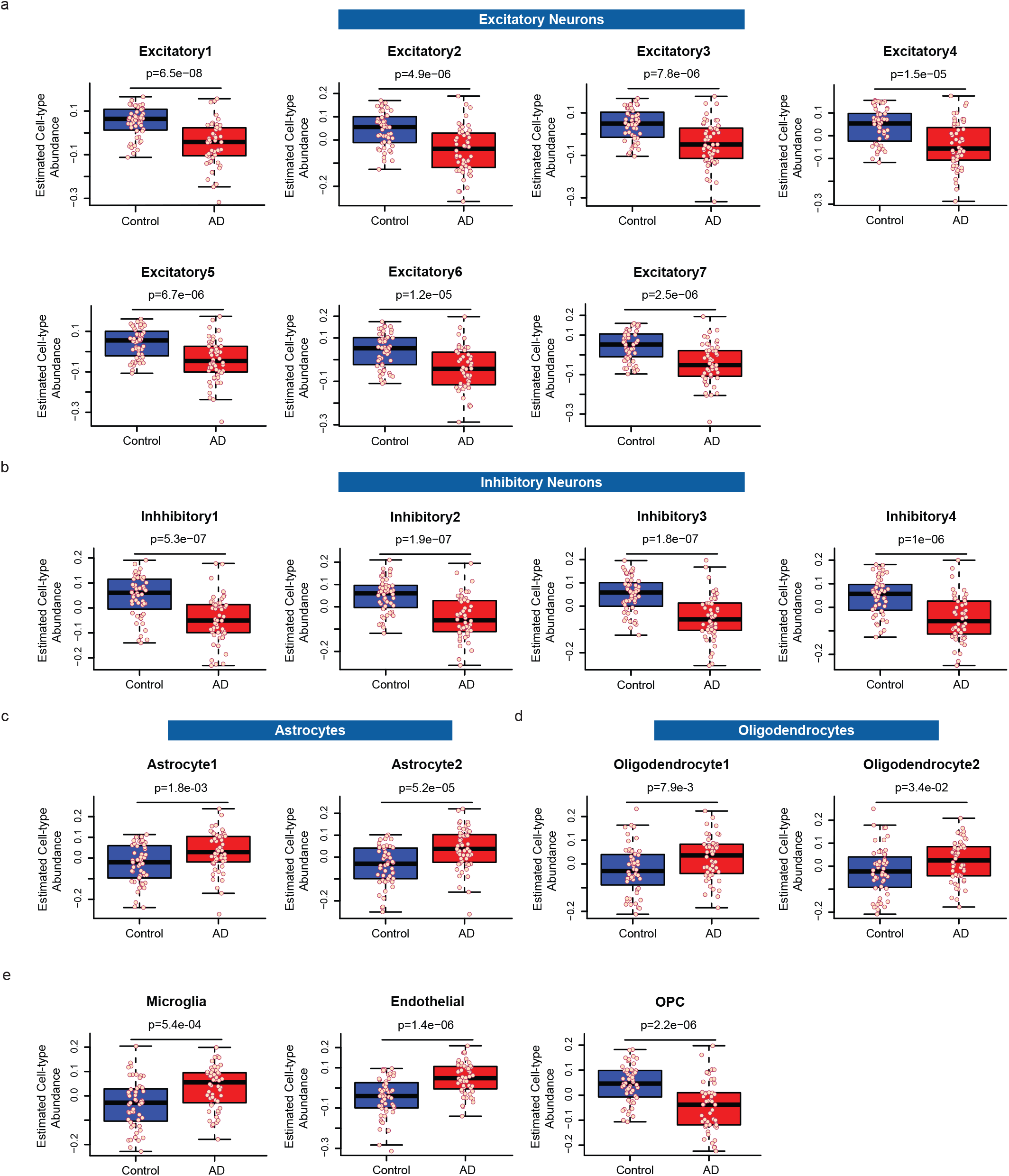
Cell-type proportion changes in AD brain (**a-e**) Boxplots depicting estimated cell-type abundance with diagnosis for excitatory neuron clusters (**a**), inhibitory neuron clusters (**b**), astrocyte clusters (**c**), oligodendrocyte clusters (**d**), and microglia, endothelial, and OPC clusters (**e**).

### Identification of disease-relevant, co-expressed transcriptional changes

Consensus weighted gene co-expression network analysis (cWGCNA)^29, 30^ is a widely used systems biology method to describe correlation patterns among genes across different samples. In order to discover consensus modules conserved across more than a thousand AD samples from different studies, we performed cWGCNA on the three bulk tissue RNA-seq datasets—Mayo, ROSMAP, and MSSM (Supplemental Figure 5a). Since these samples were from different brain regions, we aimed to identify shared and distinct transcriptional signatures across different brain regions in AD. We discovered 10 consensus modules significantly correlated with disease diagnosis—4 negatively (CM7, CM1, CM10, and CM12) and 6 positively correlated (CM16, CM5, CM4, CM8, CM9, and CM23; Figure 5a, Supplemental Table 4). We classified the consensus modules as neuronal or non-neuronal based on enrichment of cell-type markers and additionally confirmed our classification by integrating our snRNA-seq results (Figure 5b). Further, we wanted to better understand these modules by examining the relationships between them, identifying modules strongly negatively correlated, like CM1 and CM23 (Figure 5c).

**Figure 5.**
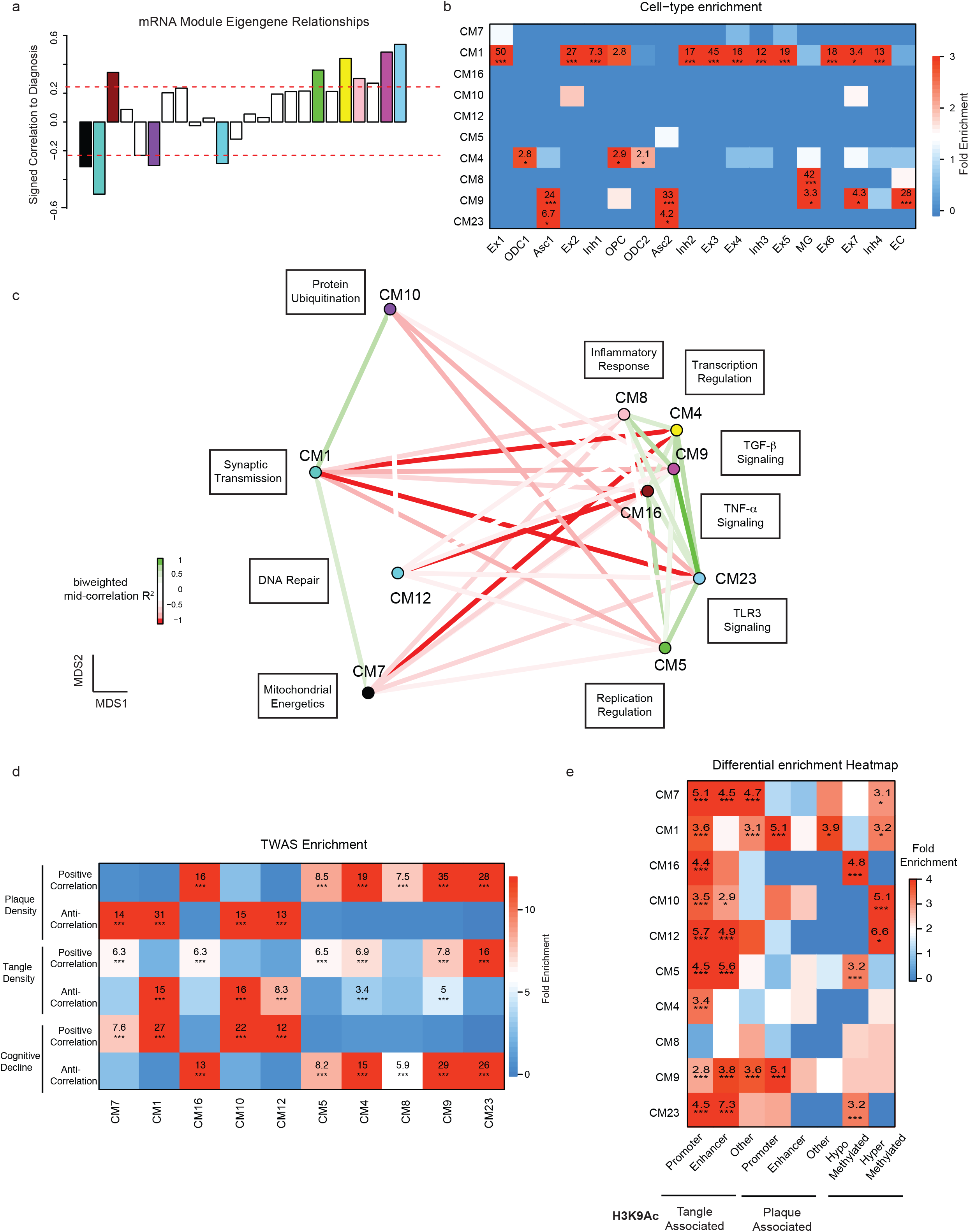
Consensus transcriptomic analysis identifies disease-relevant modules (**a**) Signed correlation of mRNA module eigengenes with diagnosis. (**b**) Enrichment of consensus modules in cell-type clusters identified by snRNA-seq. (**c**) Multidimensional scaling plot demonstrates relationships between consensus modules significantly correlated with diagnosis. Modules are also annotated with summarizing terms. (**d**) Heat map indicating enrichment of consensus modules with genes identified through transcriptome-wide association analysis (TWAS).^31^ TWAS genes were measured in relation to plaque and tangle density, as well as clinical dementia rating (cognitive decline). (**e**) Heat map showing enrichment of consensus modules with H3K9 acetylation marks, correlated with tangle and plaque burden, in promoter, enhancer or other genomic regions and AD-associated hypermethylated and hypomethylated DNA regions.

We then sought to understand the regulatory landscape of these AD-associated modules and integrated multiple layers of data to gain novel insights. We performed transcriptome-wide association studies (TWAS) to identify genes with cis-regulated expression associated with pathological and clinical traits of AD.^31^ We used plaque and tangle density as quantitative measures of the pathological changes in AD along with clinical dementia rating (CDR) scores to find significant transcriptome-wide associations. We found that downregulated consensus modules—CM1, CM7, CM10 and CM12—were enriched in TWAS genes anti-correlated with plaque and tangle density, meaning that these modules are downregulated as plaque and tangle density increases (Figure 5d). In addition, these modules were enriched in TWAS genes positively correlated with cognitive decline. Conversely, AD upregulated modules—CM4, CM16, and CM23, for example—were enriched in TWAS genes positively correlated with plaque and tangle density but negatively correlated with cognitive decline. Next, we leveraged histone H3 lysine 9 acetylation (H3K9ac) marks, which are found near active transcription start sites and enhancers,^32^ to study our consensus modules at the epigenetic level. We found that H3K9ac marks significantly associated with either tangles or Aβ plaques were enriched in all AD-associated modules with the notable exception of the glial module CM8 (Figure 5e). However, this enrichment pattern was different from AD-associated DNA methylation.^33, 34^ AD-associated hypo-methylated DNA regions were enriched in upregulated modules CM5, CM16 and CM23, whereas hyper-methylated DNA regions were enriched in all downregulated modules.

Additionally, similar to the cell-type proportion changes we found in AD, we saw that our modules had stronger correlations with disease diagnosis and the corresponding changes in module eigengene were more pronounced in datasets from the temporal cortex in comparison to the frontal cortex (Figures 6-7, Supplemental Figures 5-6). We then wanted to examine the AD-associated modules more closely and used experimentally derived databases of human protein-protein interactions (PPI) from Inweb^35^ and Biogrid^36^ to guide our attention to important genes. We created integrated co-expression-PPI networks based on the expectation that a subset of the edges defined by co-expression are due to PPI;^37–39^ such interactions functionally annotate the networks’ edges and provide orthogonal module validation with independent data.^40^ In these integrated networks, the edges between genes (nodes) represent both gene co-expression and PPI (Figure 6 a, e, i), permitting us to focus on hub genes observed at both the RNA and protein level.

**Figure 6.**
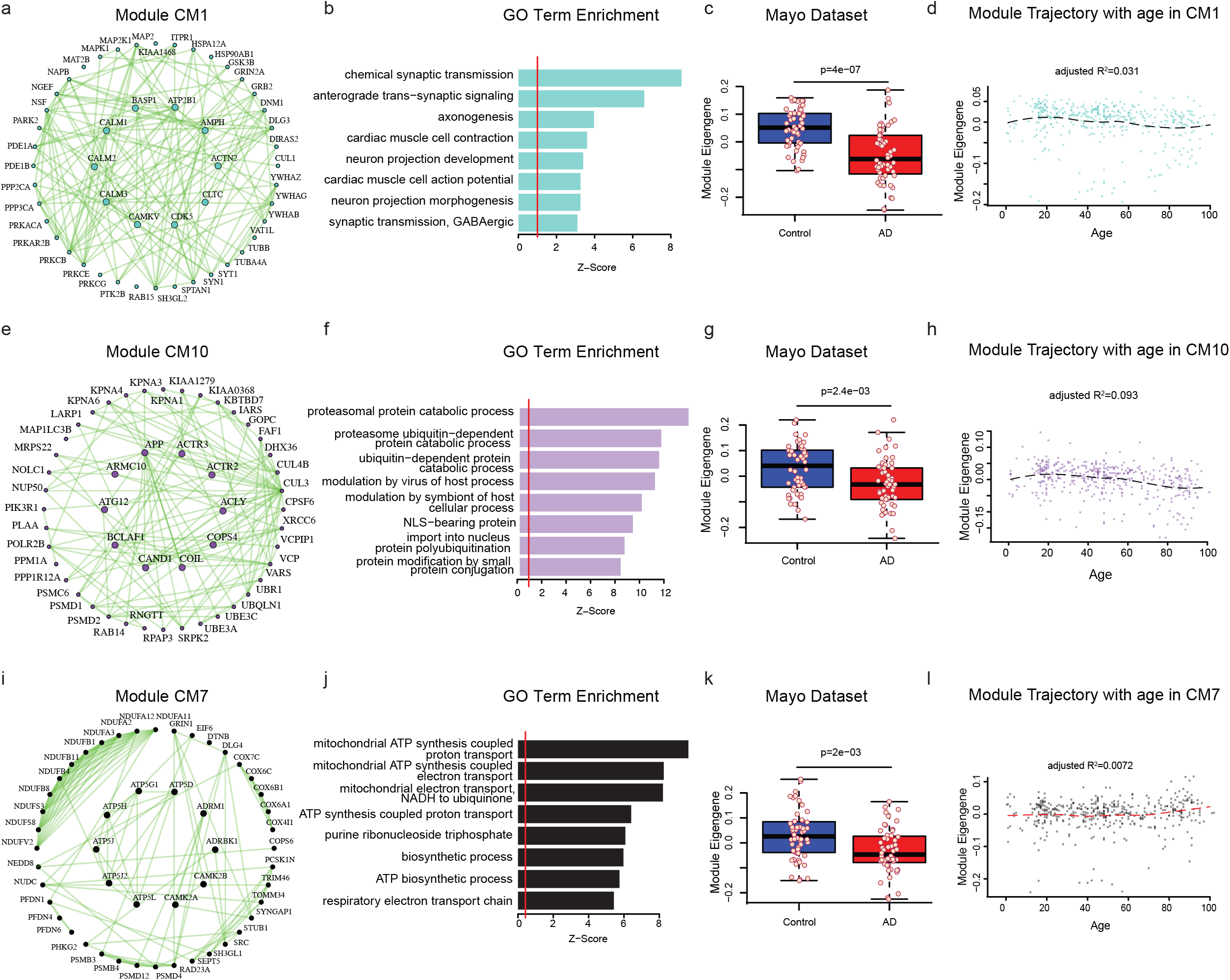
Consensus transcriptomics analysis identifies neuronal changes (**a-d**) Co-expression plot (**a**), gene ontology term enrichment (**b**), and module eigengene trajectory with diagnosis (**c**) and with age (**d**) for neuronal consensus module CM1. (**e-h**) Co-expression plot (**e**), gene ontology term enrichment (**f**), and module eigengene trajectory with diagnosis (**g**) and with age (**h**) for neuronal consensus module CM10. (**i-l**) Co-expression plot (**i**), gene ontology term enrichment (**j**), and module eigengene trajectory with diagnosis (**k**) and with age (**l**) for neuronal consensus module CM7.

**Figure 7.**
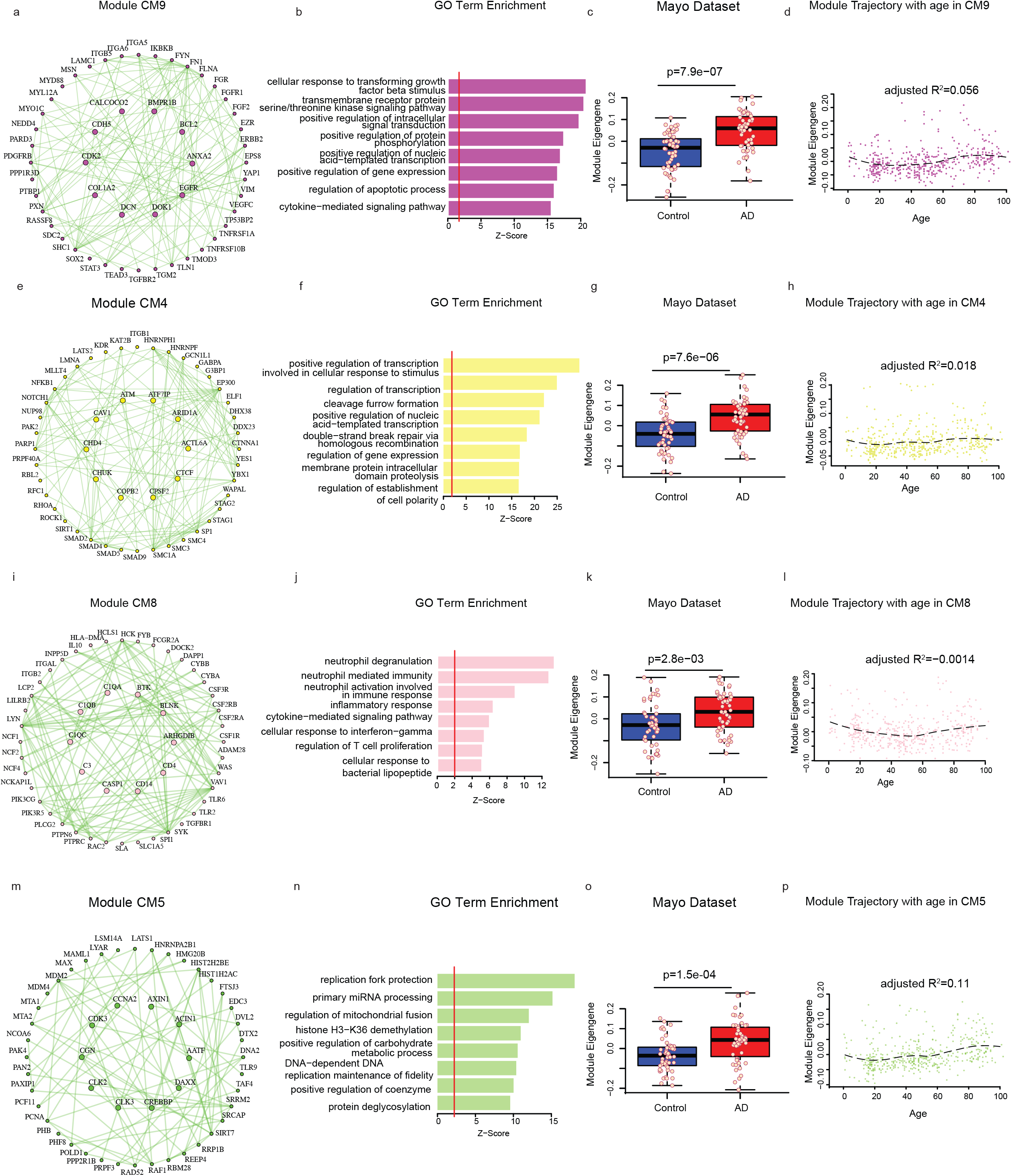
Consensus transcriptomics analysis identifies non-neuronal changes (**a-d**) Co-expression plot (**a**), gene ontology term enrichment (**b**), and module eigengene trajectory with diagnosis (**c**) and with age (**d**) for non-neuronal consensus module CM9. (**e, f, g, h**) Co-expression plot (**e**), gene ontology term enrichment (**f**), and module eigengene trajectory with diagnosis (**g**) and with age (**h**) for non-neuronal consensus module CM4. (**i-l**) Co-expression plot (**i**), gene ontology term enrichment (**j**), and module eigengene trajectory with diagnosis (**k**) and with age (**l**) for non-neuronal consensus module CM8. (**m-p**) Co-expression plot (**m**), gene ontology term enrichment (**n**), and module eigengene trajectory with diagnosis (**o**) and with age (**p**) for non-neuronal consensus module CM5.

### Neuron-specific co-expression changes in AD

Consistent with the neuronal dysfunction and neuronal loss seen in AD, consensus modules enriched in neuronal markers, such as CM1, CM10, and C7, were downregulated in AD, and these modules support the cellular changes previously identified in AD neurons. For example, CM1’s hub genes (CALM1, CALM2, CALM3, CAMKV, CLTC, and ATP2B1) are indicative of calcium signaling dysregulation, providing evidence for the calcium hypothesis of AD^41^ (Figure 6a, Supplemental Table 4), and GO analysis of CM1 demonstrated enrichment of terms like synaptic transmission and axonogenesis (Figure 6b). Moreover, one of CM10’s hub genes is APP, classically studied in AD because it transcribes the precursor of amyloid beta (Figure 6e). Notably CM10 is enriched in GO terms related to proteasomal degradation and ubiquitination (Figure 6f), and CM10’s hub gene CAND1 regulates neddylation.^42^ Neddylation has been found dysregulated in AD and may impact ubiquitination.^42^ Further, mitochondrial dysfunction and oxidative stress have been studied in neurodegeneration,^43, 44^ and specifically, mitochondria in the synapse have been found affected in AD.^45^ CM7’s hub genes include ATP synthase subunits (ATP5G1, ATP5D, ATP5L, etc.), consistent with CM7’s enrichment of GO terms related to mitochondrial function and ATP synthesis (Figure 6i-j).

In addition, we analyzed the neuronal consensus modules’ trajectories with age and with disease. We found the module eigengenes declined with disease but had no correlation with age, confirming that they were associated not with normal human aging, but instead with disease state (Figure 6).

### Glial-specific co-expression changes in AD

Glial-specific consensus modules CM9, CM4, CM8, and CM5 were positively correlated with diagnosis. Modules CM9 and CM4 highlight some of the immune-related pathways recently implicated in AD, such as TGFβ/BMP, JAK/STAT, and MAPK signaling.^46, 47^ We found CM9 was highly enriched in the astrocyte and endothelial cell-type clusters from our snRNA-seq (Figure 5b). Members of this module included genes involved in BMP (BMPR1B), JAK/STAT (STAT3), and MAPK signaling (EGFR, FGFR1, FGF2; Figure 7a), and GO analysis indicated enrichment of terms related to positive regulation of signaling pathways (Figure 7b). Further, CM4’s members were transcription factors in TGFβ signaling (SMAD2, SMAD4, etc.), and GO analysis likewise demonstrated enrichment of terms related to transcriptional regulation (Figure 7e-f).

In the same line, we identified CM8 as highly enriched in the microglia cell-type cluster (Figure 5b). Its hub genes (C1QA, C3, CASP1, CD14, etc.) and GO term enrichment of immune-related mechanisms further support a critical role of neuroinflammation in AD (Figure 7i-j). Moreover, Liddelow et al. demonstrated that astrocytes can become deleterious (A1) due to microglial secreted cytokines—one of which is C1Q.^48^ On the other hand, CM5’s GO term enrichment and genes (HNRNPA2B1, PAN2, CLK2, etc.) were associated with microRNA (miRNA) processing (Figure 7m-n). While miRNAs have been primarily studied in the context of development, miRNAs have been more recently demonstrated to play roles in the immune system and neurodegeneration.^49, 50^

We also inspected these consensus modules’ trajectories with age and with disease and found the module eigengenes increased with disease but again had no correlation with age, similar to the neuronal consensus modules (Figure 7). The remaining modules can be found in Supplemental Figure 6.

### Assessment of genetic risk within consensus modules

GWAS allows the simultaneous evaluation of thousands of genetic variants without any prior assumptions of the causal changes. Hence, we decided to examine our modules using the MAGMA pipeline, which controls for confounds like gene length. Each gene was assigned a score based on the best p-value of a single nucleotide polymorphism from an AD GWAS study,^7^ and MAGMA p-value < 0.05 was used as the cut off to include a gene as an AD candidate. We found that AD candidate genes were enriched in the microglial module CM8, further asserting a key role of the immune system in AD (Figure 8a, Supplemental Table 5). Several candidate genes, including CSF2RB, MS4A6A, CD33, SPI1 and CD14, were mapped in CM8, and these have been previously implicated in AD.^51–54^ In addition, our findings were consistent with a previous study^55^ where they reported AD risk genes in a microglial module. We also validated our results in data from genome-wide gene-based association analysis (GWGAS), which employed by Jansen et al., used MAGMA on a much larger cohort comprising of 71,880 AD cases and 383,378 controls.^31^ Similar to our GWAS enrichment scores, we found that GWGAS genes were enriched in CM8 (Figure 8b). Moreover, we found that CM8 was enriched in AD-associated brain expression quantitative trait loci (eQTLs, see Methods), while another glial module, CM9, was enriched in AD-associated brain methylation QTLs (mQTLs, see Methods).

**Figure 8.**
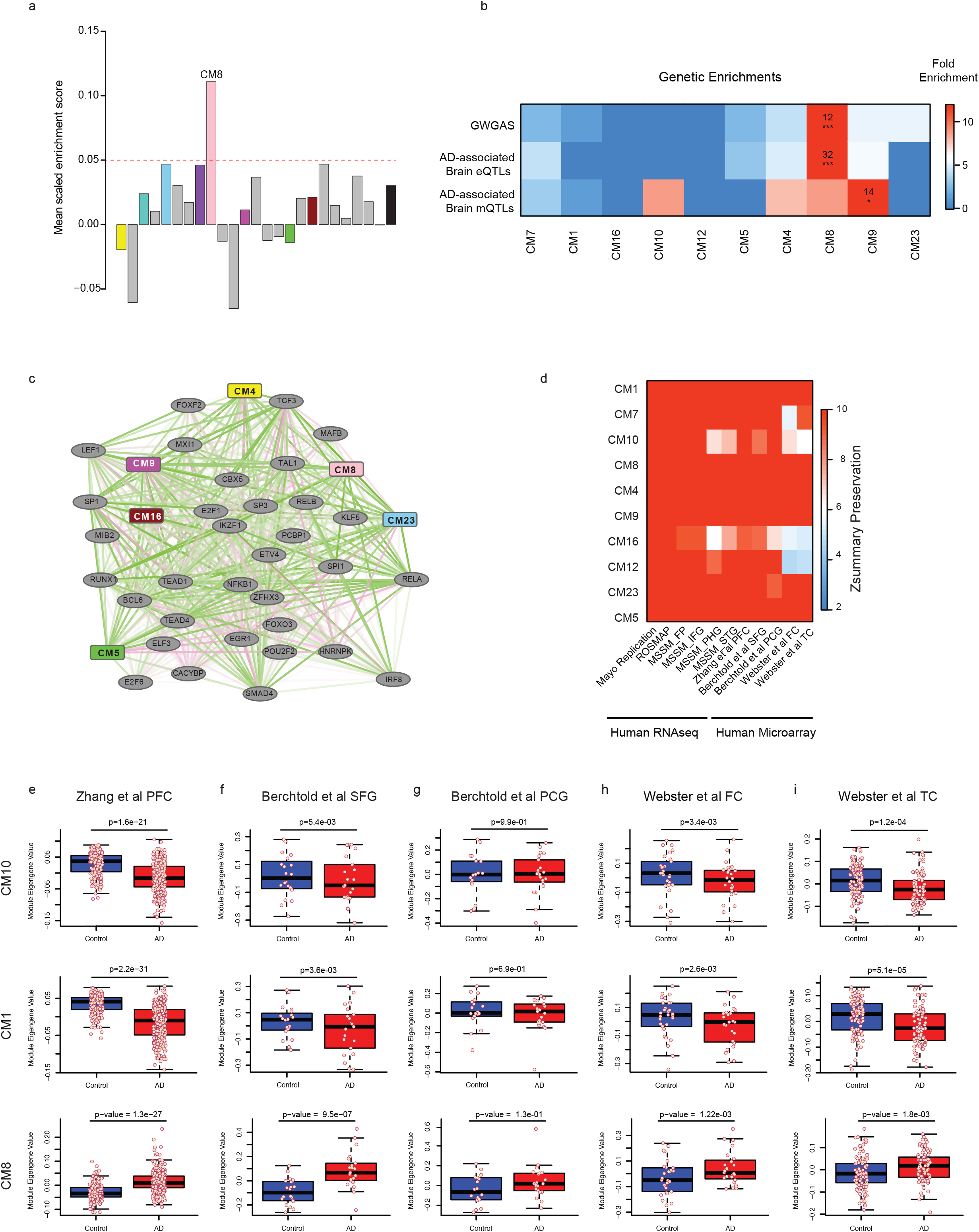
Genetic enrichment and validation of findings (**a**) Mean scaled enrichment of AD genome-wide association study (GWAS) hits from Lambert et al.^7^ in consensus modules. (**b**) Heat map showing enrichment of consensus modules in genome-wide gene-based association analysis (GWGAS) hits and AD-associated brain expression and methylation quantitative trait loci (eQTLs and mQTLs). (**c**) Co-expression plot demonstrating demonstrates relationships between non-neuronal consensus modules and transcription factors. (**d**) Heat map indicating module preservation in other human RNA-seq and microarray datasets. (**e-i**) Boxplot showing module eigengene trajectory with diagnosis of consensus modules CM10, CM8 and CM1 in datasets Zhang et al. PFC^55^ (**e**), Berchtold et al. superior frontal gyrus (SFG**, f)** and post-central gyrus^65^ (PCG, **g**), and Webster et al. FC (**h**) and TC^66^ (**i**). The upper and lower lines represent the 75^th^ and 25^th^ percentile respectively, and the central line represents the median.

### Identification of transcription factors regulating consensus modules

Network analysis identifies co-expressed modules that are likely co-regulated.^38, 56, 57^ With this in mind, we performed transcription factor binding site enrichment analysis (TFBS, see Methods) to identify transcription factors (TFs) that may be regulating disease-associated modules. This method has been successfully used previously to accurately predict TFs.^38^ TFBS revealed that master TF—RELA, otherwise known as p65, appears to control both neuronal and non-neuronal modules (Figure 8c, Supplemental Figure 7a). It becomes further evident that NFκB signaling may be mediating transcriptional changes in AD since we found transcription factors NKFB1 and RELB associated with non-neuronal modules (Figure 8c). Previous studies have reported increased NFκB activation in AD and pointed to a relationship with Aβ pathology.^58–61^ Moreover, Withaferin A has been identified as an inhibitor of NFκB signaling^62, 63^ and has shown therapeutic potential to protect against Aβ toxicity.^64^

### Validation of consensus modules in other datasets

Since modules can be affected by non-biological variables like differences in sample preparation, we needed to authenticate the existence of our modules of interest in other datasets. In order to do this, we utilized module preservation analysis, which helps us quantify the features of a particular module that are preserved across multiple networks (see Methods). We found that our consensus modules were highly preserved across published human RNA-seq and microarray datasets,^55, 65, 66^ indicated by Z-summary preservation (Figure 8d, Supplemental Table 6).

Moreover, we examined the module eigengene trajectories of modules—CM10, CM1 and CM8—in these datasets and found comparable changes with diagnosis (Figure 8e-i). As expected and also in line with our findings, we observed changes were notably greater in temporal cortex than frontal cortex, again reflecting AD’s progressive atrophy of different brain regions^1^ (Supplemental Figures 7-8). These results altogether support the robustness of our findings.

## Discussion

Our multi-scale study across different brain regions using high-dimensional datasets provides a comprehensive picture of the genome-wide transcriptomic alterations occurring in AD and begins to examine the associated gene regulatory programs. While GWAS along with studies in familial AD cases have identified several causal genetic variants associated with the disease, ^6, 7^ these findings have not been sufficient for fully understanding neurodegeneration since it has been difficult to ascertain the functional significance of these genetic variants. We hypothesized that AD culminates from multiple genetic variants, not only making the molecular mechanisms of AD complex, but also resulting in the great phenotypic variability. Thus, we aimed to identify transcriptional changes that are conserved across multiple large cohorts, utilizing integrative genomics and systems biology approaches to yield novel insights into disease mechanisms. We hope that our findings will pave way for accelerated drug development, a critical need in AD, since our study has identified robust co-expressed modules and their hubs, which can be targets for novel therapeutics.

Bulk tissue transcriptomic approaches from various model systems of disease, including postmortem human brain samples, provide a direct approach to understand disease processes in affected disease-relevant tissues. Analyzing multiple cohorts across different cortical regions has its advantage as we gain more power to accurately quantify gene expression changes across hundreds of samples. However, the diversity of brain banks coupled with differences in RNA-seq library preparation and quantification methods poses additional challenges in analyzing all the samples together. Here we have used a standardized data analysis approach to uniformly process (data QC and alignment, see Methods) all the samples and quantify gene expression changes across multiple brain regions. Using uniformly processed samples, we were able to identify differentially expressed genes in both the protein-coding and non-coding transcriptome and validate these findings bioinformatically in multiple datasets. Differential expression analysis quickly highlighted an important limitation of bulk transcriptomic approaches—they are inadequate for understanding cell-type proportion changes in AD. These bulk tissue RNA-seq approaches also do not address cell-type heterogeneity-associated changes that occur in the disease.

To address these issues, we generated single-cell transcriptomic profiles on more than 27,000 nuclei from control human prefrontal cortex are generated, and the gene expression signatures were used to cluster nuclei in an unbiased manner. We found that our single-cell approach is robust in identifying all major neural cell types, including multiple clusters for excitatory and inhibitory neurons, oligodendrocytes, and astrocytes. In addition, we used a novel algorithm to estimate cell-type abundance in bulk RNA-seq data with a single-cell transcriptomic dataset. Using this algorithm, we were able to calculate cell-type specific markers with high intramodular connectivity (kME). Subsequently, genes with high kME values (kME>0.7) were used to calculate the first principal component (PC1) in bulk RNA-seq datasets. With this approach, we were able to show that the abundance of several excitatory and inhibitory neuronal clusters is decreased in AD cases, whereas the abundance of microglial and astrocyte clusters is increased in AD as compared to control.

In order to put bulk transcriptomic data in a systems-level framework, we performed cWGCNA on the ROSMAP, Mayo and MSSM datasets. cWGCNA has been successfully used in the past to identify transcriptomic signatures conserved in disease and to find meaningful biological relationships^29, 30^. cWGCNA on AD cohorts identified robust conserved and biologically important transcriptomic alterations across multiple brain regions in AD. Using this approach, we found neuronal modules, downregulated in AD, that are related to synaptic transmission, calcium signaling (CM1), ubiquitination (CM10) and mitochondrial dysfunction (CM7). Additionally, we found glial modules associated with immune response (CM8) and JAK/STAT and MAPK signaling (CM9 and CM4) upregulated in AD.

We further studied the consensus modules by integrating TWAS genes and H3K9ac marks associated with tangle and plaque density. Downregulated modules were clearly positively correlated with CDR scoring but negatively correlated with pathological measurements, while upregulated modules were the opposite. Interestingly, we noticed the upregulated glial module CM8 was not enriched in any H3K9ac marks. Moreover, one of the biggest challenges in co-expression analysis is to find robust co-expressed modules that are altered in different cohorts and datasets. In order to tackle this problem, we used module preservation analysis, which is a powerful tool to find conserved modules in different datasets. Through module preservation analysis, we found that our AD-associated consensus modules are robustly preserved in multiple published AD datasets from different brain regions.^55, 65, 66^

Since aging is the most important non-genetic risk factor for AD, we wanted to investigate if the AD-associated consensus modules are altered during normal human aging. Normal aging has been shown to alter glial and neuronal gene-expression changes in the brain;^67^ however, we hypothesized that these changes are distinct from pathological changes occurring in neurodegenerative diseases. Identification of neuronal and glial changes, which do not alter in normal human aging but are only associated with AD, can provide us with mechanistic insights into disease biology. We calculated synthetic module eigengenes from AD-associated consensus modules in a North American Brain Expression Consortium (NABEC) dataset^67^ to show that the AD-associated modules are not significantly co-related during normal aging. The NABEC dataset we used was comprised of brain gene expression data from hundreds of normal human patients between the age of 16-100 years.^67^ This finding in combination with the robust reproducibility of our modules in published AD datasets^55, 65, 66^ demonstrated that we have identified disease-specific transcriptomic signatures in AD.

A key issue in any analysis of gene expression in disease is that changes in gene expression may either be a cause or a consequence of the disorder being studied. In order to address this potential issue, we ascertained specific enrichment of causal genetic factors in disease-related co-expression modules.^38, 55, 56, 68^ We found significant enrichment of AD GWAS common genetic risk signals in the immune-related module CM8, which is enriched in microglial marker genes. These data support the potential causal role of these modules containing known risk genes and suggest causal pathways in AD consistent with previous reports of glial enrichment of AD GWAS data.^55, 69^ Our work provides the framework for disease-associated gene regulatory networks in AD and provides the foundation for future work in drug discovery.

## Acknowledgements

For ROSMAP dataset, the study data were provided by the Rush Alzheimer’s Disease Center, Rush University Medical Center, Chicago. Data collection was supported through funding by NIA grants P30AG10161, R01AG15819, R01AG17917, R01AG30146, R01AG36836, U01AG32984, U01AG46152, the Illinois Department of Public Health, and the Translational Genomics Research Institute. The results published here are in whole or in part based on data obtained from the AMP-AD Knowledge Portal (doi:10.7303/syn2580853). For Mayo dataset, the study data were provided by the following sources: The Mayo Clinic Alzheimer’s Disease Genetic Studies, led by Dr. Nilufer Taner and Dr. Steven G. Younkin, Mayo Clinic, Jacksonville, FL using samples from the Mayo Clinic Study of Aging, the Mayo Clinic Alzheimer’s Disease Research Center, and the Mayo Clinic Brain Bank. Data collection was supported through funding by NIA grants P50 AG016574, R01 AG032990, U01 AG046139, R01 AG018023, U01 AG006576, U01 AG006786, R01 AG025711, R01 AG017216, R01 AG003949, NINDS grant R01 NS080820, CurePSP Foundation, and support from Mayo Foundation. Study data includes samples collected through the Sun Health Research Institute Brain and Body Donation Program of Sun City, Arizona. The Brain and Body Donation Program is supported by the National Institute of Neurological Disorders and Stroke (U24 NS072026 National Brain and Tissue Resource for Parkinson’s Disease and Related Disorders), the National Institute on Aging (P30 AG19610 Arizona Alzheimer’s Disease Core Center), the Arizona Department of Health Services (contract 211002, Arizona Alzheimer’s Research Center), the Arizona Biomedical Research Commission (contracts 4001, 0011, 05-901 and 1001 to the Arizona Parkinson’s Disease Consortium) and the Michael J. Fox Foundation for Parkinson’s Research. The results published here are in whole or in part based on data obtained from the AMP-AD Knowledge Portal (doi:10.7303/syn2580853). For MSSM dataset, the data were generated from postmortem brain tissue collected through the Mount Sinai VA Medical Center Brain Bank and were provided by Dr. Eric Schadt from Mount Sinai School of Medicine. The results published here are in whole or in part based on data obtained from the AMP-AD Knowledge Portal (doi:10.7303/syn2580853). In addition, we are grateful to Thanneer Perumal and Ben Logsdon at Sage bionetworks for processing the fastq and bam files used in the analysis.

## Data and code availability

RNA-seq data and details can be found on https://www.synapse.org using corresponding synapse (syn) IDs-Mayo RNA-seq (syn5550404), MSBB (syn3159438) and ROSMAP (syn3219045). Single-cell data can be found on synapse website using synID# syn18915937. Additionally we have created shiny apps to easily visualize gene, network-level and single-cell data which can be accessed through our lab website at https://swaruplab.bio.uci.edu/consensusAD. Code for consensus WGCNA can be found here: https://github.com/vswarup/ConsensusWGCNA

## Methods

### Description of human samples and RNA-seq data

For single-cell analysis control human pre-frontal cortex brain samples were obtained from UCI MIND’s Alzheimer’s disease research center (ADRC) brain bank. These samples were obtained under UCI institutional review board (IRB). Post-mortem human brain tissues were obtained from three cohorts – Mayo study, Mount Sinai Brain Bank (MSBB) study and Religious Orders Study and Memory and Aging Project (ROSMAP) study. All the human samples were obtained under their respective institutional review board. For details on human samples used in this study and their availability, please see Supplemental Table 1. RNA-seq data and details can be found on synapse.org website using corresponding synapse (syn) IDs-Mayo RNA-seq (syn5550404), MSBB (syn3159438) and ROSMAP (syn3219045).

### Other published datasets used in the study

For normal human aging, we used North American Brain Expression Consortium (NABEC) data ranging from 16 to 101 years old from frontal cortex.^67, 70, 71^ The gene-expression arrays are available from Gene expression omnibus (GEO) - GSE36192. Published AD microarray datasets were also downloaded from GEO including Zhang et al prefrontal cortex data^55^ (GSE44770), Berchtold et al. frontal and temporal cortex data^65^ (GSE48350) and Webster et al.^66^ (GSE15222).

### RNA-seq data processing

We performed standard alignment, QC and gene-expression analysis on each of the three datasets (ROSMAP, Mayo and MSSM) to ensure uniform processing of data. All samples were aligned to the human genome (GRCh38, GENCODE v25 gene model) using rnaSTAR aligner. Counts were obtained using rnaSTAR quantification and conditional quantile normalized (CQN) FPKM data was used for downstream analysis.

### Differential gene expression analysis

Differential gene expression analysis was done using TPM data and effects from biological covariates like diagnosis, age, gender, post-mortem interval (PMI), and technical variables like batch (depending on the cohort), RNA integrity number (RIN) and sequencing biases were assed using a linear regression model. The final model used was implemented in R is as follows:

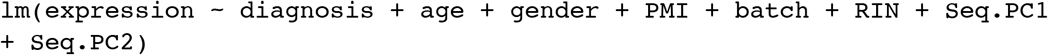

where Seq.PC1 and Seq.PC2 are sequencing PCs obtained from aggregating sequencing metrics obtained from Picard tools.

### Consensus WGCNA

Consensus weighted co-expression network analysis (cWGCNA) was performed with ROSMAP, Mayo and MSSM datasets using WGCNA package in R.^29^ We employed a signed cWGCNA approach by calculating component-wise values for topological overlap for individual brain banks. First, bi-weighted mid-correlations were calculated for all pairs of genes, and then a signed similarity matrix was created. In the signed network, the similarity between genes reflects the sign of the correlation of their expression profiles. The signed similarity matrix was then raised to power β (β=12) to emphasize strong correlations and reduce the emphasis of weak correlations on an exponential scale. The resulting adjacency matrix was then transformed into a topological overlap matrix as described elsewhere.^38^ Modules were defined using specific module cutting parameters which included minimum module size of 100 genes, deepSplit=4 and threshold of correlation = 0.2. Modules with correlation greater than 0.8 were merged together. We used first principal component of the module, called module eigengene, to correlate with diagnosis and other variables. Hub genes were defined using intra-modular connectivity (kME) parameter of the WGCNA package. Gene-set enrichment analysis was done using enrichR package.^72^

### Module Preservation Analysis

To understand the functional relationship of modules and its network properties in orthogonal datasets, we performed module preservation analysis. Module definitions from the cWGCNA analysis was used and Z-statistics were calculated using the module preservation function in WGCNA package in R.^73^

### Genetic enrichment with eQTLs, mQTLs and GWGAS

Brain specific eQTLs were generated from databases including the CommonMind Consortium Portal, AMP-AD, BRAINEAC and xQTL Serve. A false discovery rate of 0.05 was used to define significant eQTL associations. eQTLs which overlapped with AD associated SNPs from either Lambert et al and Jansen et al GWAS datasets^6, 7^ were annotated as AD-associated eQTLs. Similarly, methylation QTLs (mQTLs) defined in AMP-AD, and CommonMind Consortium Portal were annotated as AD-associated mQTLS if they overlapped with AD GWAS data. We also used genome-wide gene-based association analysis (GWGAS) data using Multi-marker Analysis of GenoMic Annotation (MAGMA).^74^ GWAS enrichment analysis in modules was done using MAGMA pipeline. Briefly, GWAS summary statistics was obtained for AD from IGAP website.^7^ Using MAGMA approach,^74^ we assigned each gene a score based on the best P value of a SNP in a given GWAS study within 20kB of the gene, and then set a P value cut-off at 0.05 to define the gene as included in the common variant set related to AD. We performed enrichment analysis with logistic regression controlling for gene length, LD blocks and other biases.^38^

### TWAS Enrichment Analysis

We performed AD TWAS using the FUSION package (http://gusevlab.org/projects/fusion/)^31^ with gene-expression measured in ROSMAP PFC data and GWAS summary statistics from AD GWAS.^7^ We chose the best model of accuracy in the FUSION package (best cis-eQTL, best linear unbiased predictor, Bayesian sparse linear mixed model (BSLMM),^75^ Elastic-net regression, LASSO regression) and used for downstream association analyses. TWAS association statistics were Bonferroni corrected per GWAS, gene and intron separately.

Association for TWAS was assessed for plaque and tangle densities and genome-wide significant genes were used for enrichment analysis. Similarly, clinical dementia rating (CDR) reported in ROSMAP dataset was used for association with cognitive decline.

### Single-cell Library Preparation

Five control human brain samples were processed for nuclei isolation following modified version of the published protocol.^23^ All procedures were carried on ice. ∼ 50 mg of the brain sample was homogenized in 500 μl chilled Nuclei EZ lysis buffer (Sigma-Aldrich), using a motorized tissue grinder and disposable pellet pestles (Fisher Scientific) by stroking ∼10-20 times. The homogenate was then incubated for 5 minutes after adding 1 ml of the Nuclei EZ lysis buffer. Homogenate was then filtered using a 70 μm MACS strainer (Miltenyi Biotec). The flow through was centrifuged at 500g for 5 minutes at 4°C and the supernatant was carefully removed. The nuclei pellet was then re-suspended in another 1.5 ml of Nuclei EZ lysis buffer and incubated for 5 minutes. The cell suspension was again centrifuged (500g,5 minutes, 4°C) and the supernatant was removed. 500 μl of Nuclei wash and re-suspension buffer (NWR buffer: 1xPBS (Fisher Scientific), 1%BSA (Sigma-Aldrich) and 0.2U/ μl RNase inhibitor (New England Bio Labs)) was then added and incubated without re-suspending, to allow buffer interchange. After the incubation, 1ml of NWR buffer was added and the nuclei was re-suspended. The resuspension was centrifuged (500g,5 minutes, 4°C) and the supernatant was removed. The nuclei pellet was then gently resuspended in 1.4 ml of NWR buffer, centrifuged (500g, 5 minutes, 4°C), removed supernatant and was gently re-suspended in another 500 μl of NWR buffer. All nuclei were collected by washing the walls of the centrifuge tube. Nuclei re-suspension was then filtered using 40-μm Flowmi cell strainer (Bel-Art). Number of nuclei was then counted using an automated cell counter (Bio-Rad). Samples were directly sorted by FACS using DAPI positive gates at the UCI Immunology FACS core in the 10X buffer and libraries were prepared using Single-cell RNA-seq library kit (v3, 10X Genomics). Samples were pooled and sequenced at an average read-depth of 100,000 reads/cell.

### Single-cell Clustering and analysis

Raw sequencing Illumina bcl2 files were demultiplexed using 10X Genomics Cell Ranger software. Further analysis was performed in Seurat V3 package in R.^76^ We used sctransform function in Seurat to normalize the data and top 75 PCs were used for clustering and visualization using Fourier transformation t-SNE (FItsne^77^). Differential expression between clusters were performed using MAST package in Seurat.^78^

### Cell-type composition analysis

Using the most variable genes in a cluster, we performed WGCNA analysis and determined the intramodular connectivity (kME) of each gene. kME values are correlation of first principal components of a module/cluster to the expression of that gene. Thus, higher kME values can often explain most of the variance in the data because of their high correlation to PC1. Using Fisher’s transformation, we converted kME values to specificity scores (Z-scores). We then used the top one percentile of genes by their kME values and calculated their module eigengene (ME) in respective bulk RNA-seq dataset. These ME values were then plotted to understand the cell-composition changes in control and AD samples.

### Transcription factor binding site enrichment (TFBS)

TFBS enrichment was done using pipeline published elsewhere.^56^ Briefly the computational pipeline identifies the transcription factors (in the promoter region) in the given gene list using experimentally validated transcription factors (TFs). The pipeline used TF position-weighted matric from JASPAR and TRANSFAC databases as well as includes experimental associated datasets obtained from ENCODE^79^ and ChEA.^80^ The pipeline uses clover algorithm and corrects for background using human chromosome 22.

*Supplemental Table 1* – Metadata for postmortem brain samples used for bulk tissue RNA-seq and single-nuclei RNA-seq which includes Mayo cohort (Discovery and Replication set), ROSMAP, Mt Sinai MSSM, single-nuclei RNA-seq data (UCI).

*Supplemental Table 2* – Differential Expression analysis showing log2 fold change and associated p-values obtained using linear regression model on Mayo discovery set samples.

*Supplemental Table 3* – Single-nuclei RNA-seq of postmortem human brain – Table showing original clusters based on Seurat analysis as well as cell ID cluster based on marker gene expression. Differential expression (DE) analysis using MAST package showing top DE genes.

*Supplemental Table 4* – Consensus WGCNA analysis describing all the module colors and module labels as well as intramodular connectivity (kME) values.

*Supplemental Table 5* – MAGMA GWAS enrichment analysis results in each module showing fold enrichment and p-values.

*Supplemental Table 6* – Module preservation z-summary and Bonferroni corrected p-value, where consensus WGCNA modules were used as reference module.

---

**Supplemental Figure 1.**
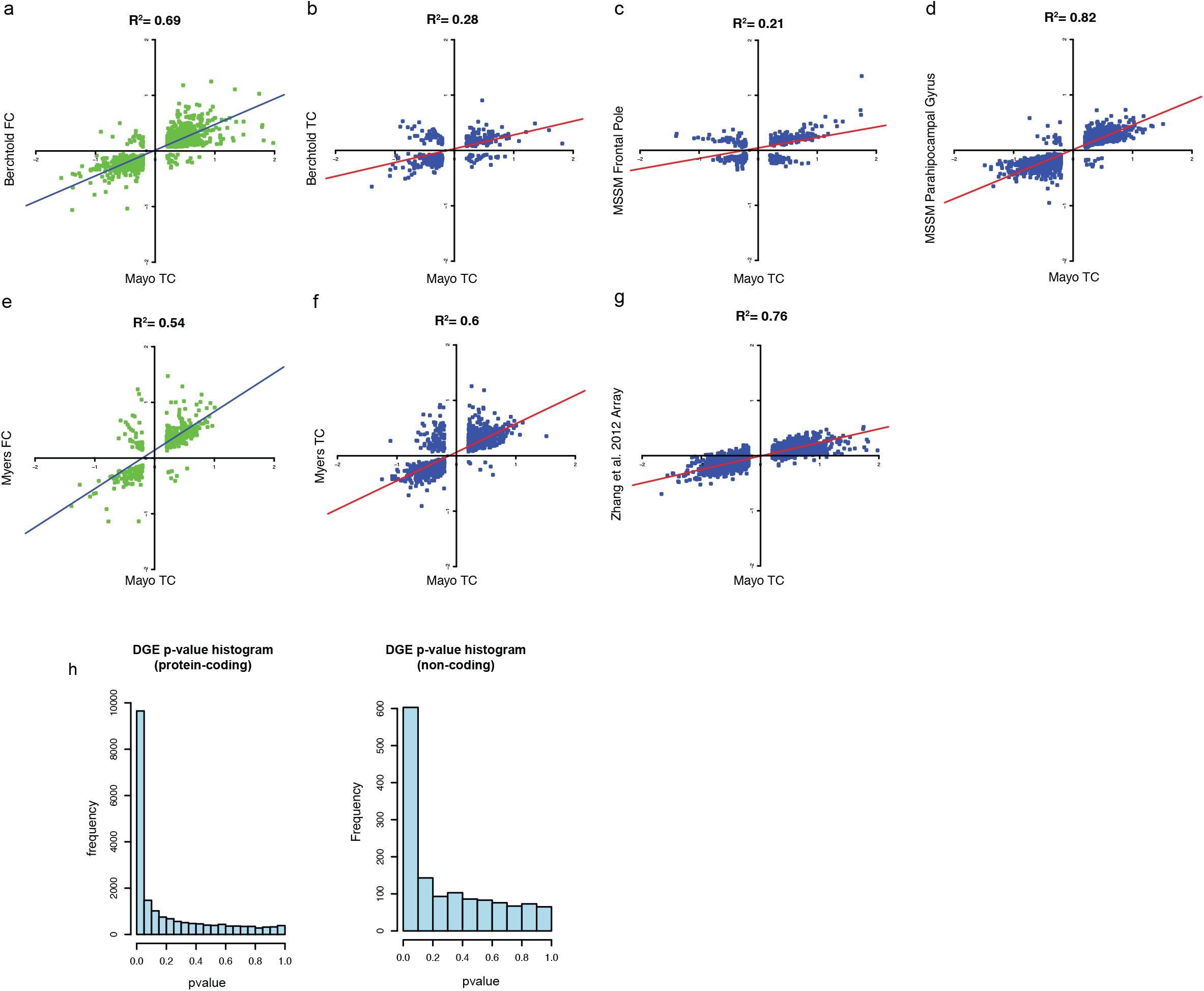
Validation of differential expression signature in ROSMAP and MSSM datasets. (**a-g**) Correlation of differentially expressed genes from Mayo temporal cortex (TC) dataset with Berchtold frontal cortex (FC, **a**), Berchtold TC (**b**), MSSM frontal pole (**c**), MSSM para-hippocampal gyrus (**d**), Webster FC (**e**), Webster TC (**f**), and Zhang et al., FC (**g**). (**h**) Histogram of p-values for differentially expressed protein-coding (left) and non-coding (right) genes.

**Supplemental Figure 2.**
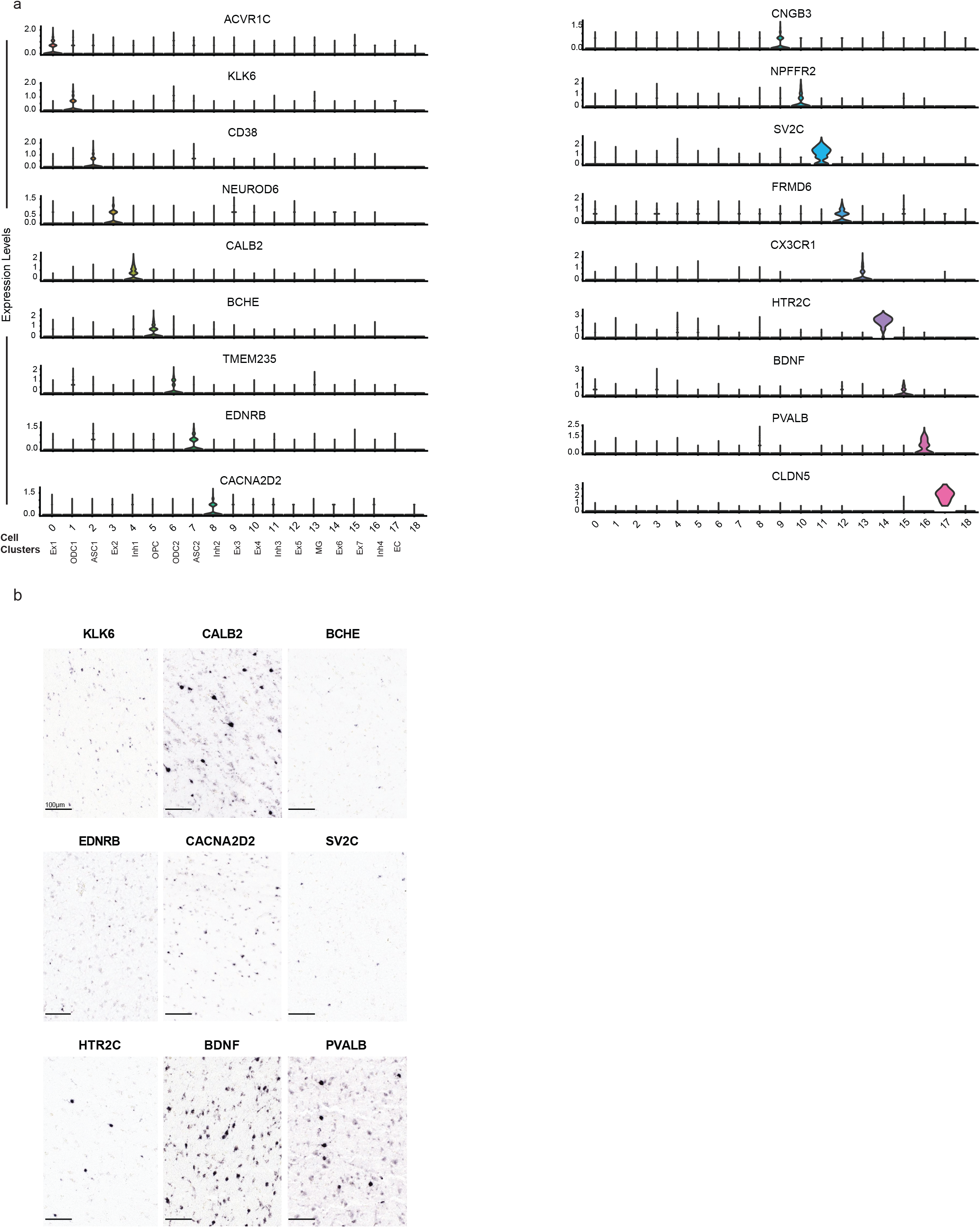
Single-cell cluster ISH and marker gene expression. (**a**) Violin plots of differentially expressed cell-type markers in each cell-type cluster identified by single-cell RNA-seq. (**b**) RNA ISH of a subset of the differentially expressed cell-type markers in human cortical tissue. Images from the Allen Human Brain Atlas.

**Supplemental Figure 3.**
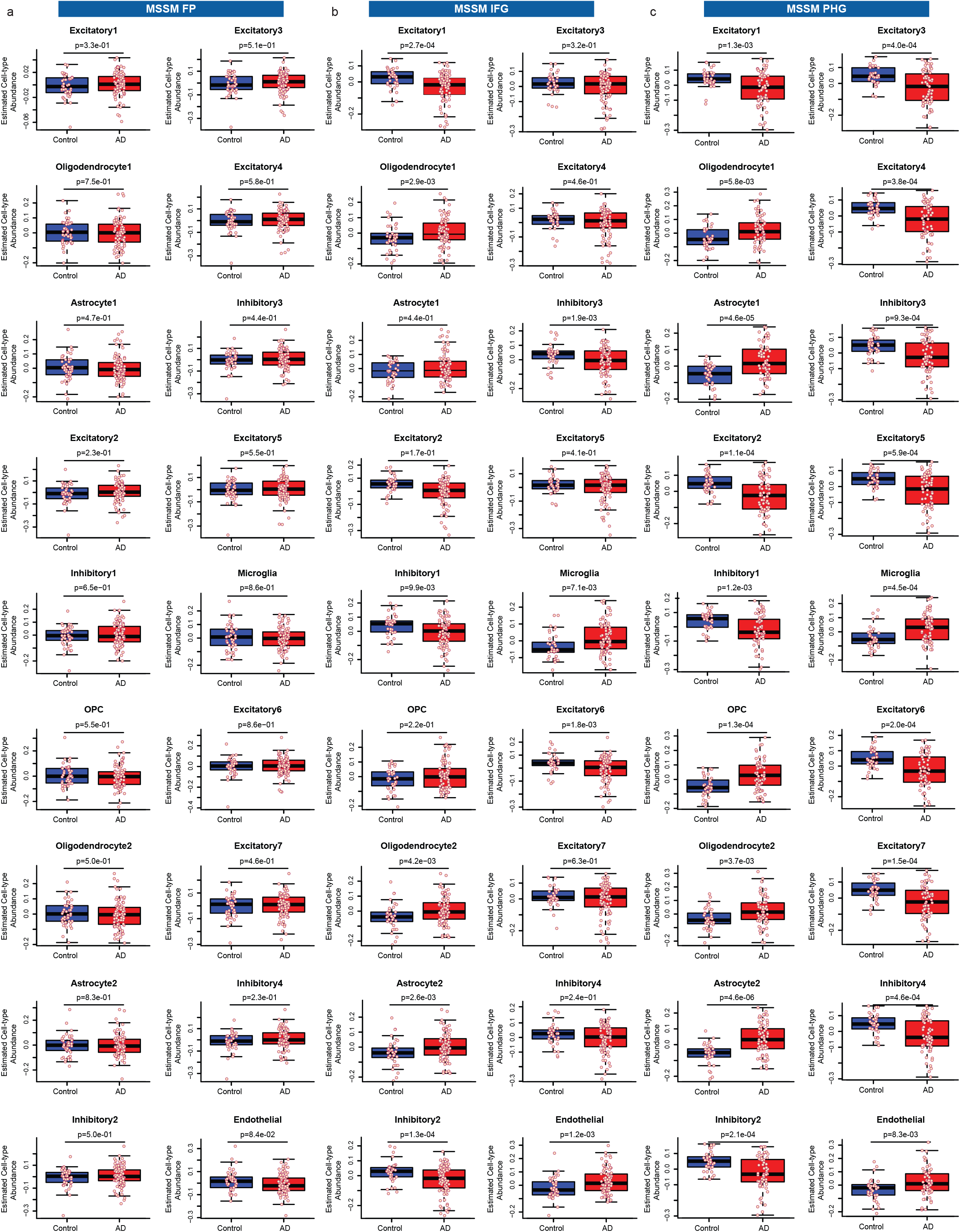
Cell-type proportion changes in MSSM datasets. (**a-c**) Boxplots depicting estimated cell-type abundance with diagnosis by cell-type cluster in MSSM RNA-seq dataset from FP (**a**), IFG (**b**), and PHG (**c**).

**Supplemental Figure 4.**
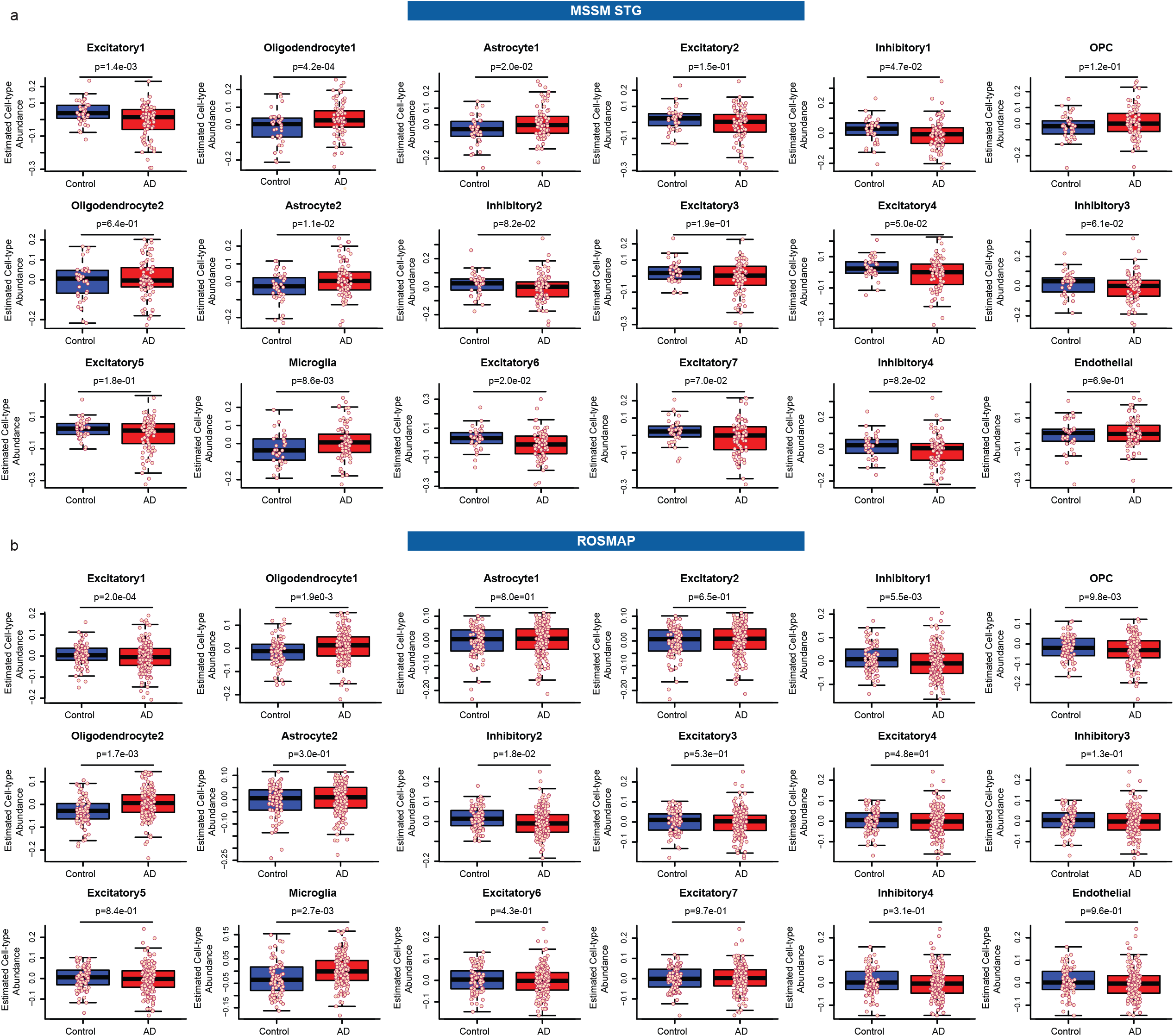
Cell-type proportion changes in MSSM and ROSMAP datasets. (**a-b**) Boxplots depicting estimated cell-type abundance with diagnosis by cell-type cluster from RNA-seq dataset from MSSM STG (**a**) and ROSMAP PFC (**b**).

**Supplemental Figure 5.**
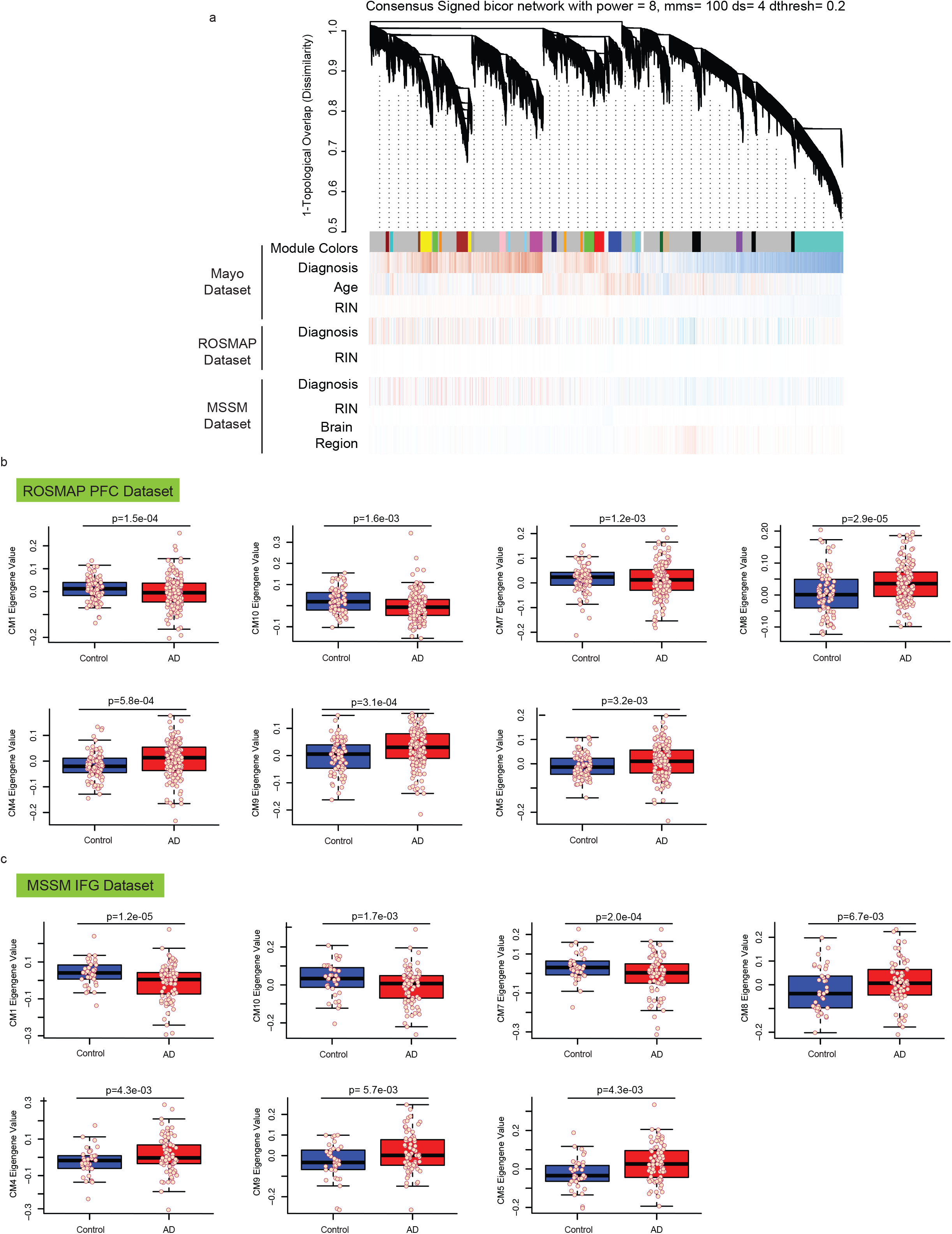
Consensus WGCNA analysis in ROSMAP and MSSM datasets. (**a**) Consensus WGCNA dendrogram showing consensus modules from Mayo, ROSMAP, and MSSM datasets. Red indicates positive correlation of covariate (diagnosis, age, RIN) with gene expression, and blue is negative correlation. (**b-c**) Boxplot showing module eigengene trajectory with diagnosis for neuronal and non-neuronal consensus modules in ROSMAP PFC (**b**) and MSSM IFG (**c**) RNA-seq datasets.

**Supplemental Figure 6.**
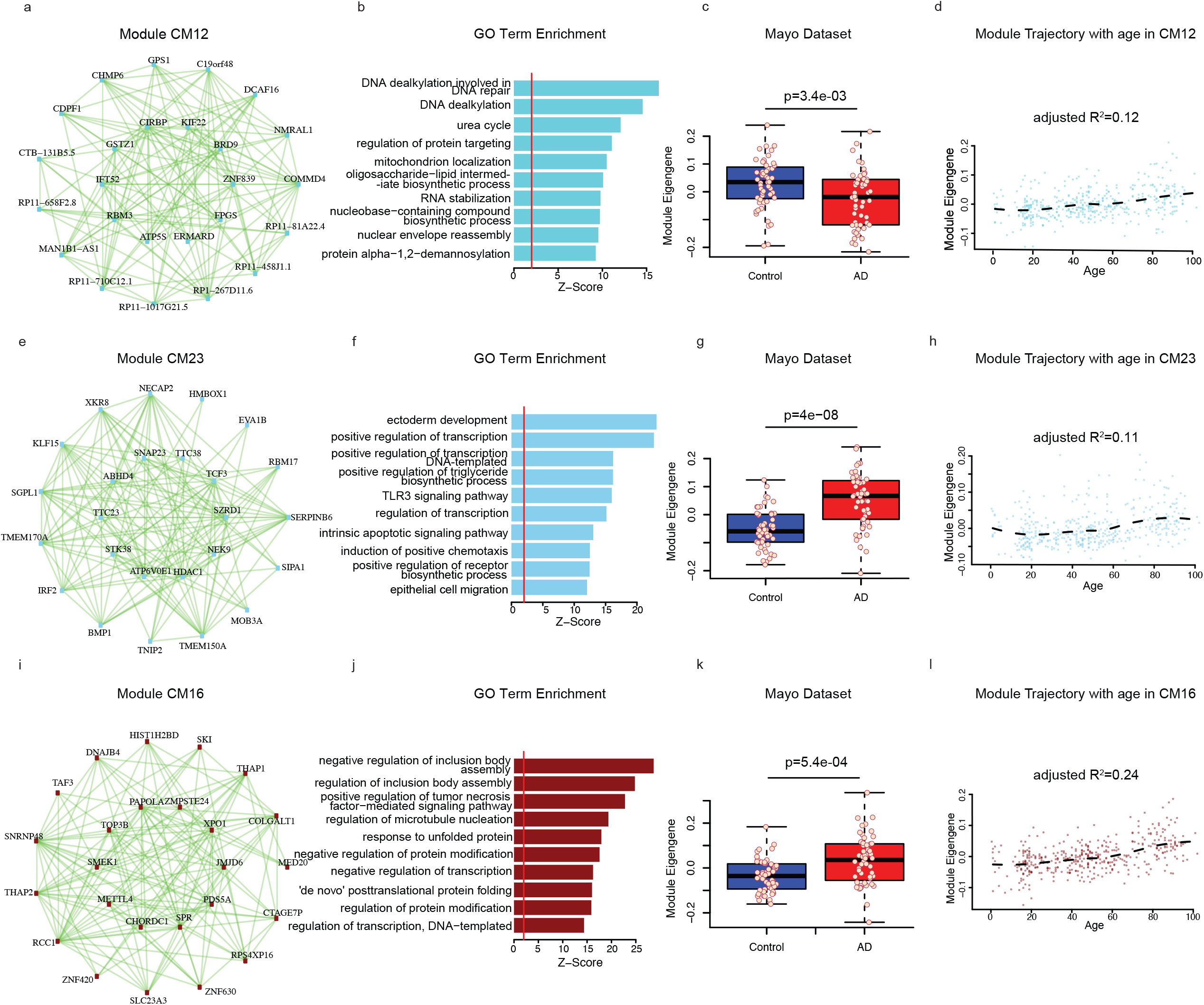
ME trajectory, pathway analysis and module plots of CM12, CM23 and CM16 modules. (**a-d**) Co-expression plot (**a**), gene ontology term enrichment (**b**), and module eigengene trajectory with diagnosis (**c**) and with age (**d**) for consensus module CM12. (**e-h**) Co-expression plot (**e**), gene ontology term enrichment (**f**), and module eigengene trajectory with diagnosis (**g**) and with age (**h**) for consensus module CM23. (**i-l**) Co-expression plot (**i**), gene ontology term enrichment (**j**), and module eigengene trajectory with diagnosis (**k**) and with age (**l**) for consensus module CM16.

**Supplemental Figure 7.**
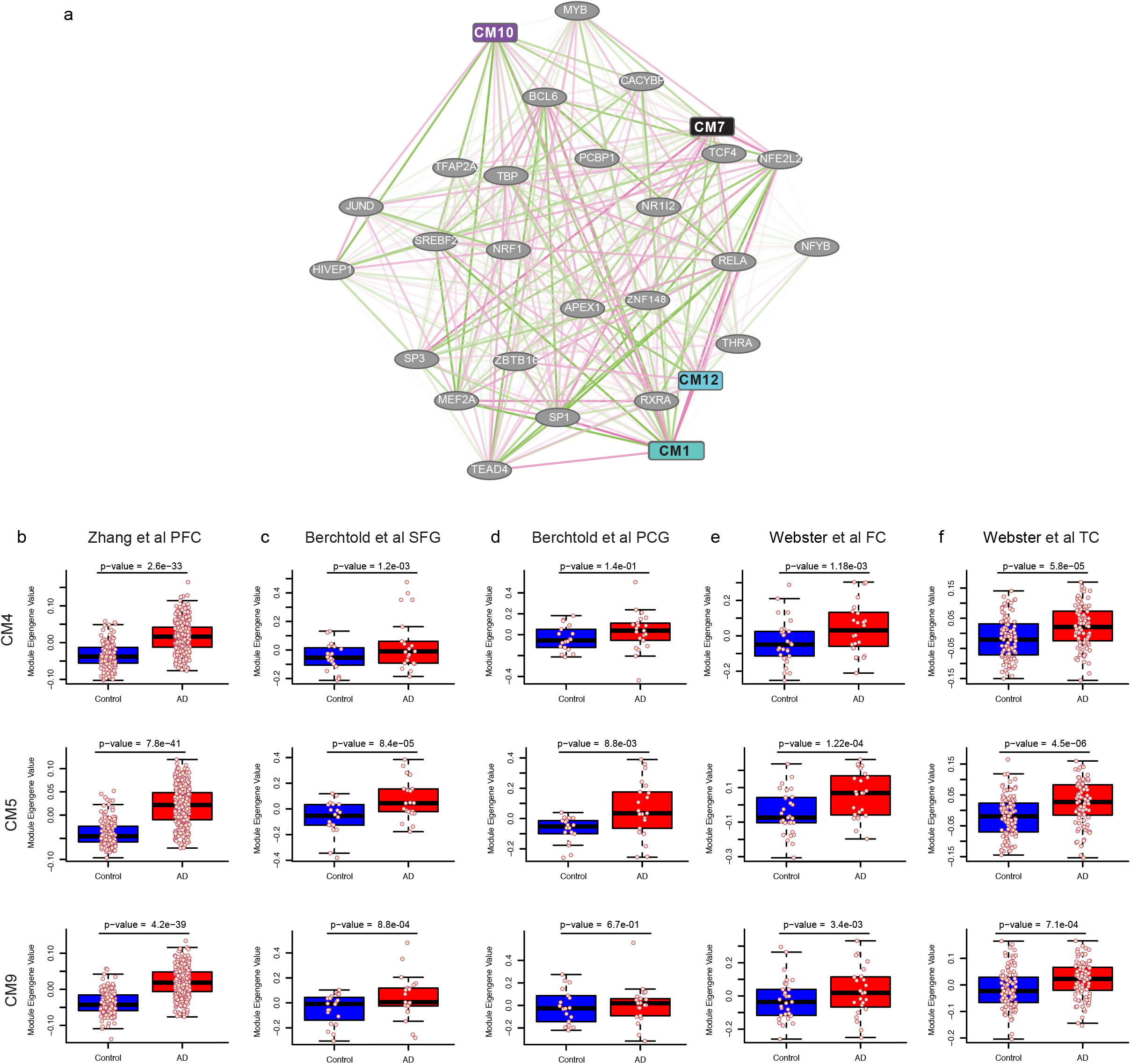
Transcription factors regulating neuronal modules and ME trajectory in published AD datasets. (**a**) Co-expression plot demonstrating relationships between neuronal consensus modules and transcription factors. (**b-f**) Boxplots illustrating module eigengene trajectory with diagnosis from RNA-seq dataset from Zhang et al., PFC (**b**), Berchtold et al., SFG (**c**) and PCG (**d**), and Webster et al., FC (**e**) and TC (**f**).

**Supplemental Figure 8.**
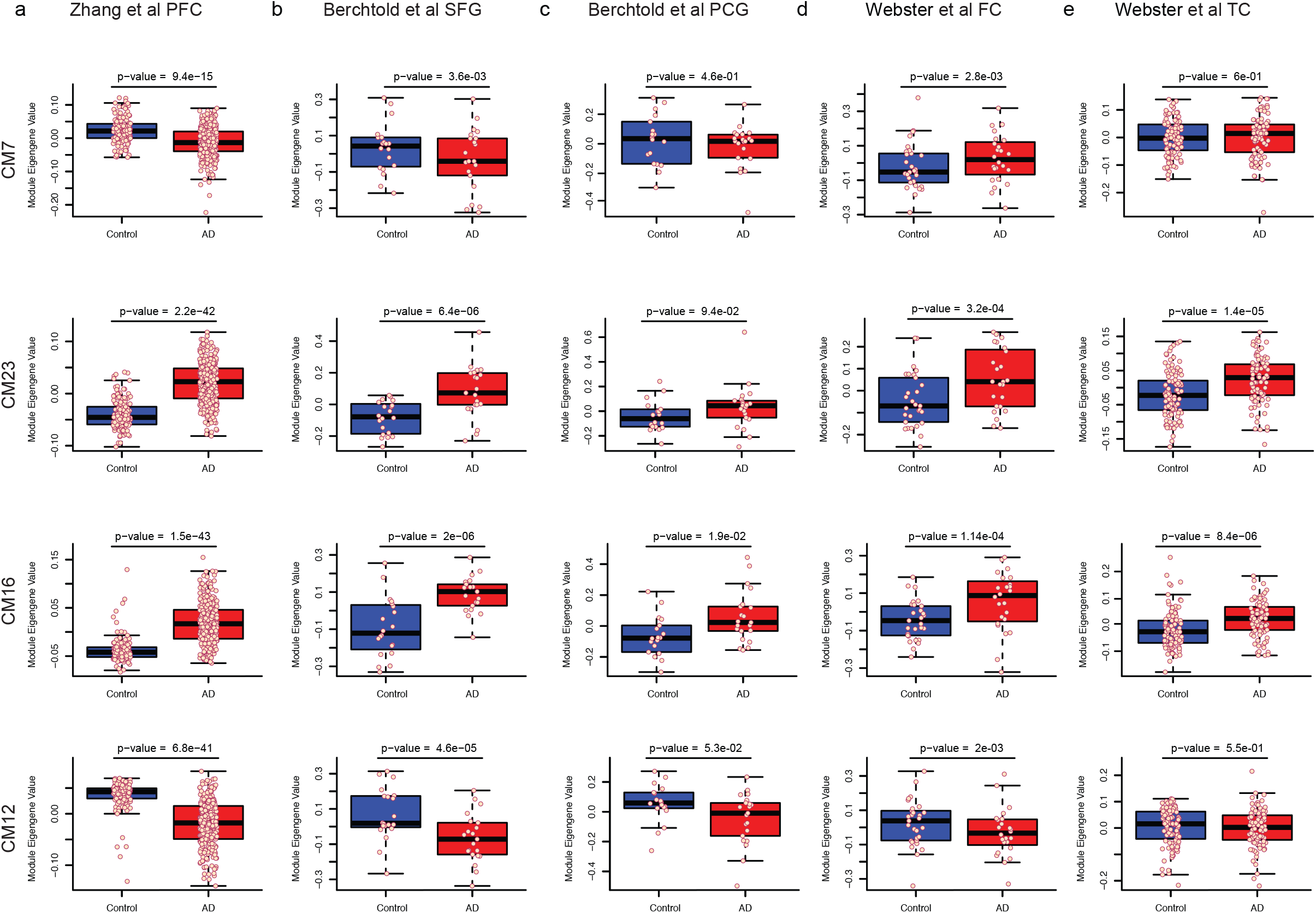
ME trajectory in published AD datasets. (**a-e**) Boxplots illustrating module eigengene trajectory with diagnosis from RNA-seq dataset from Zhang et al., PFC (**a**), Berchtold et al., SFG (**b**) and PCG (**c**), and Webster et al., FC (**d**) and TC (**e**).

